# RFX6-mediated dysregulation defines human β cell dysfunction in early type 2 diabetes

**DOI:** 10.1101/2021.12.16.466282

**Authors:** John T. Walker, Diane C. Saunders, Vivek Rai, Chunhua Dai, Peter Orchard, Alexander L. Hopkirk, Conrad V. Reihsmann, Yicheng Tao, Simin Fan, Shristi Shrestha, Arushi Varshney, Jordan J. Wright, Yasminye D. Pettway, Christa Ventresca, Samir Agarwala, Radhika Aramandla, Greg Poffenberger, Regina Jenkins, Nathaniel J. Hart, Dale L. Greiner, Leonard D. Shultz, Rita Bottino, Human Pancreas Analysis Program, Jie Liu, Stephen C.J. Parker, Alvin C. Powers, Marcela Brissova

**Affiliations:** Department of Molecular Physiology and Biophysics, Vanderbilt University School of Medicine, Nashville, Tennessee, 37232, USA; Division of Diabetes, Endocrinology and Metabolism, Department of Medicine, Vanderbilt University Medical Center, Nashville, Tennessee, 37232, USA; Department of Computational Medicine and Bioinformatics, University of Michigan, Ann Arbor, Michigan, 48109, USA; Department of Human Genetics, University of Michigan, Ann Arbor, Michigan, 48109, USA; Department of Molecular Medicine, Diabetes Center of Excellence, University of Massachusetts Medical School, Worcester, Massachusetts, 01655, USA; The Jackson Laboratory, Bar Harbor, Maine, 04609, USA; Imagine Pharma, Devon, Pennsylvania, 19333, USA; Institute of Cellular Therapeutics, Allegheny-Singer Research Institute, Allegheny Health Network, Pittsburgh, Pennsylvania, 15212, USA; Human Islet Research Network (RRID:SCR_014393); Department of Biostatistics, University of Michigan, Ann Arbor, Michigan, 48109, USA; VA Tennessee Valley Healthcare System, Nashville, Tennessee, 37212, USA

## Abstract

A hallmark of type 2 diabetes (T2D), a major cause of world-wide morbidity and mortality, is dysfunction of insulin-producing pancreatic islet β cells^1–3^. T2D genome-wide association studies (GWAS) have identified hundreds of signals, mostly in the non-coding genome and overlapping β cell regulatory elements, but translating these into biological mechanisms has been challenging^4–6^. To identify early disease-driving events, we performed single cell spatial proteomics, sorted cell transcriptomics, and assessed islet physiology on pancreatic tissue from short-duration T2D and control donors. Here, through integrative analyses of these diverse modalities, we show that multiple gene regulatory modules are associated with early-stage T2D β cell-intrinsic defects. One notable example is the transcription factor RFX6, which we show is a highly connected β cell hub gene that is reduced in T2D and governs a gene regulatory network associated with insulin secretion defects and T2D GWAS variants. We validated the critical role of RFX6 in β cells through direct perturbation in primary human islets followed by physiological and single nucleus multiome profiling, which showed reduced dynamic insulin secretion and large-scale changes in the β cell transcriptome and chromatin accessibility landscape. Understanding the molecular mechanisms of complex, systemic diseases necessitates integration of signals from multiple molecules, cells, organs, and individuals and thus we anticipate this approach will be a useful template to identify and validate key regulatory networks and master hub genes for other diseases or traits with GWAS data.

## INTRODUCTION

Type 2 diabetes mellitus (T2D), a metabolic disease defined by hyperglycemia, is a major cause of macro and microvascular morbidity and mortality for more than 460 million individuals worldwide^1^. Clinically heterogenous, T2D involves genetic, environmental, and physiologic components that impact multiple molecular pathways and tissues^2, 3^. Initial management frequently involves diet and lifestyle alterations but often escalates to require multiple oral or injectable medications and ultimately exogenous insulin to lower blood glucose^7, 8^. T2D is associated with obesity and age, both of which reduce peripheral tissue sensitivity to insulin; however, most individuals with insulin resistance do not develop T2D. Instead, the key defining feature of those who develop T2D is impaired insulin secretion^7, 9^. Insulin is secreted endogenously by the β cell within the pancreatic islet. In addition to β cells, the islet also contains other endocrine cells (α, δ, γ, and ε), vascular structures (endothelial cells and pericytes), and immune cells, which collectively function as a mini-organ to control glucose homeostasis in a coordinated fashion^10, 11^. While islet dysfunction is a hallmark of T2D, it remains unclear whether this is the result of an intrinsic β cell defect, a reduction in β cell number, systemic signals from altered levels of fatty acids, glucose, or lipids, or some combination of these.

T2D has a strong genetic component with more than 400 signals identified through genome-wide association studies (GWAS)^4^. Loci linked to T2D through GWAS are enriched in β cell-specific open chromatin regions, suggesting impaired β cell processes as a key determinant for whether T2D develops and how quickly it progresses^5, 6^. Further, 90% of GWAS-identified single nucleotide polymorphisms (SNPs) are located in non-coding parts of the genome, and they are enriched in predicted islet enhancer regions where many likely modulate cell-specific gene expression regulatory networks by altering transcription factor binding^12–16^. How personalized genetic variation causes changes in cell-specific gene and protein expression, tissue architecture, and cellular physiology in T2D islets is not well understood.

Postulated T2D disease processes include β cell loss and/or dedifferentiation, endoplasmic reticulum (ER) stress, amyloid deposition, oxidative stress, glucotoxicity, lipotoxicity, and islet inflammation^17–20^. These processes have been primarily studied in rodent models of T2D due to difficulty in obtaining and studying human pancreatic tissue and islets. Importantly, human islets show several key differences from mouse islets, including endocrine and non-endocrine cell composition and arrangement, basal and stimulated insulin secretion, response to dyslipidemia and hyperglycemia, and expression of key islet-enriched transcription factors^21–24^, highlighting the need for studies to define initiating and sustaining mechanisms of islet dysfunction in primary human islets.

Recent advances in cadaveric pancreas procurement and processing have increased availability of human tissue for histological analysis as well as ex vivo molecular and functional profiling of islets isolated from individuals with diabetes. However, many studies utilize only tissue or islets, and further, do not differentiate study outcomes based on T2D duration. Since different stages of T2D may involve different processes, studies that combine cases with different T2D duration make it difficult to discern cellular and molecular causes from disease consequences. The association of physiological measurements with transcriptomic profiles of islet cells have begun to identify key pathways for β cell function^25, 26^, but integration of these studies with disease stage, tissue-based analyses, and genetic risk remains a challenge.

Here, we used an integrated approach to study the pancreas and isolated islets from donors with short-duration T2D and nondiabetic controls to identify disease-driving molecular defects early in the course of T2D. We analyzed islet function both ex vivo and in vivo using a transplant system and performed comprehensive transcriptional analysis by bulk RNA-sequencing (RNA-seq) of whole islets and purified β cells and α cells, correlating these profiles to functional parameters and GWAS variants using weighted gene co-expression network analysis (WGCNA). Concurrently, we described changes in the pancreatic islet microenvironment via traditional and multiplexed imaging approaches, including assessing spatial cell relationships. We found that dysfunction in short-duration T2D is defined primarily by β cell-intrinsic defects, including an RFX6-governed and GWAS-enriched transcriptional regulatory network.

## RESULTS

### Identification, collection, and processing of short-duration T2D donor pancreata

To identify early, disease-driving mechanisms in islets, we focused on short-duration T2D as defined by a combination of disease duration and treatment approach (**Fig. 1a**). Using a national network, we identified high-quality organs to ensure minimal ischemic time and consistently applied multiple tissue processing and fixation methods, including simultaneous collection of isolated islets and tissue from the same pancreas when possible. Twenty pancreata were obtained from individuals with T2D aged 37-66y (mean 52y) with T2D duration of 0-10y (mean 3.5y). Of these donors, 25% were without pharmaceutical treatment (HbA1c range 6.2-9.9; mean 7.6) and 75% were on diabetes medication, mostly oral agents (HbA1c range 6.3-11.2; mean 8.0) (**Fig. 1a**). Pancreata from nondiabetic (ND) donors (n=17) were also collected and processed for multi-modality study. Partnerships with the Integrated Islet Distribution Program (IIDP) and the Alberta Diabetes IsletCore provided access to additional islets from ND donors (n=19) to assist with matching of donor characteristics. Detailed information, including sample types and experimental usage for each case, is available in **Extended Data Table 1**. Application of multiple modalities allowed for integrative analysis of ex vivo and in vivo islet function, tissue architecture and microenvironment including spatial relationships, and cell type-specific gene expression (**Fig. 1b**).

**Figure 1.**
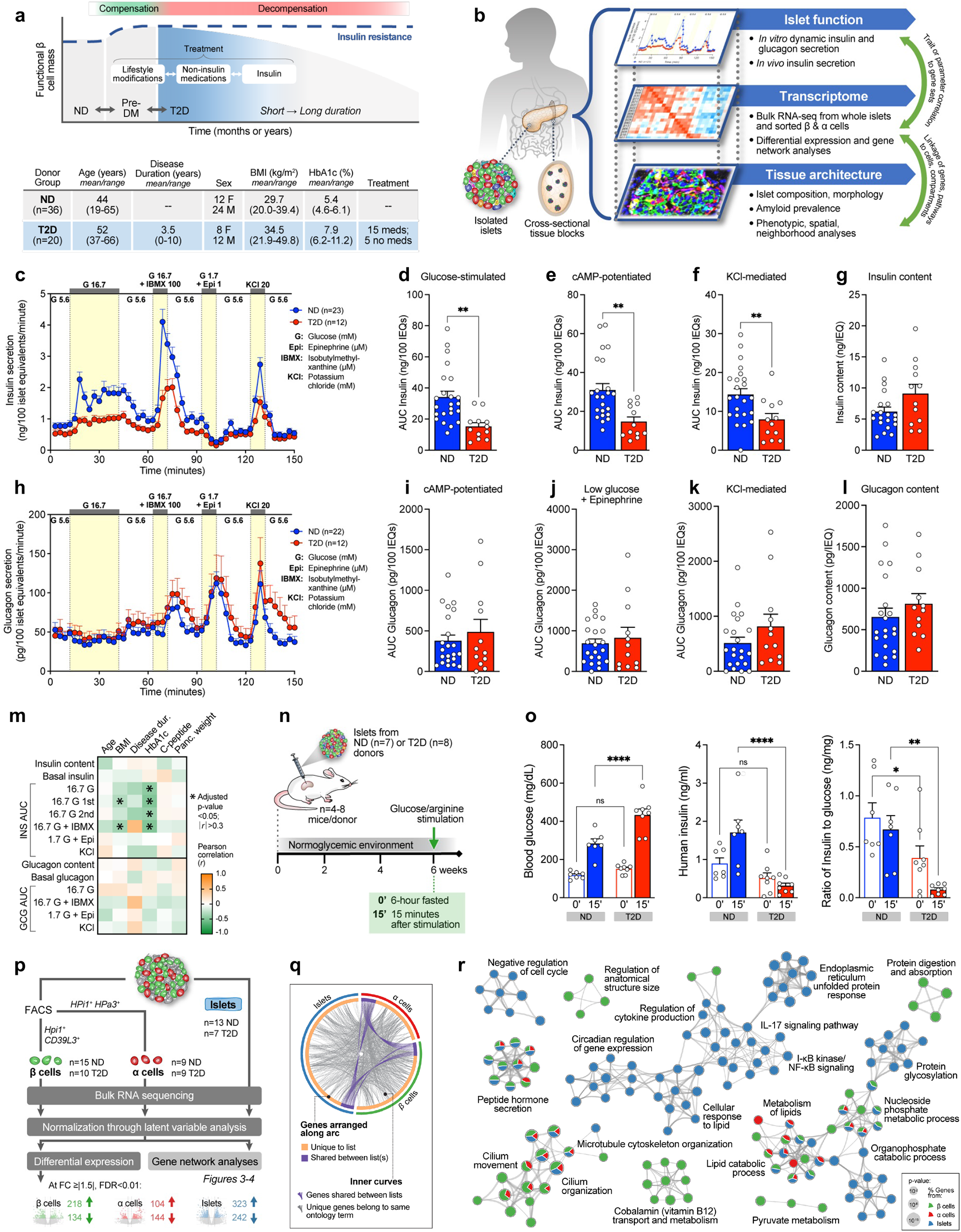
Integrated analysis of islet function, gene expression, and histology in a cohort of donors with short-duration type 2 diabetes (T2D) reveals substantially reduced stimulated insulin secretion ex vivo and in vivo despite similar insulin content and highlights dysregulated pathways in purified β and α cells as well as whole islets. **(a)** Schematic of functional β cell mass during disease progression from nondiabetic (ND) to pre-diabetes (Pre-DM) and T2D, highlighting the divergence of insulin supply and demand and escalation of treatment mirroring progressive loss of functional β cell mass. Shaded blue represents targeted disease stage in this cohort with clinical profile shown below in table. **(b)** Schematic of multimodal study of islet function, transcriptome, and tissue architecture. Coordinated study on islets and tissue from same donor allowed integration between analyses (green arrows). **(c-l)** Dynamic insulin and glucagon secretory responses measured by islet perifusion. Panels **d-f** and **i-k**: secretagogue response as area under the curve (AUC); **g, l**: hormone content normalized to islet volume. **(m)** Pearson correlation of perifusion metrics to clinical traits. **(n)** Schematic of human islet transplantation and in vivo assessment of function. **(o)** Blood glucose, human insulin levels, and human insulin:blood glucose ratio measured before and after glucose and arginine stimulation of mice with human islet grafts. Symbols show donor average. **(p)** Schematic of RNA sample collection and analysis. **(q)** Overlap of differentially expressed (DE) genes in T2D β cell (green) α cell (red), and islet (blue) samples at the level of genes (purple curves) or ontology terms (grey curves). **(r)** Metascape network showing a subset of enriched terms from DE genes. Edges denote similarity and node colors reflect contribution of sample(s). * p<0.05, ** p<0.01, *** p<0.001, **** p<0.0001 (**d-g**, **i-l**: two-tailed t-test; **o**: two-way ANOVA).

### Short-duration T2D islets show reduced stimulated insulin secretion

To investigate islet function, we assessed dynamic hormone secretion in isolated islets from age-and body mass index (BMI)-matched T2D and ND donors (**Extended Data Fig. 1a-1b**) by a standardized perifusion approach that interrogates multiple steps of the insulin secretory pathway and has been adopted by the Human Islet Phenotyping Program of the IIDP to assess over 400 human islet preparations^27^. When normalized by islet volume, stimulated insulin secretion was substantially reduced in response to high glucose, cyclic AMP (cAMP)-evoked potentiation, and potassium chloride (KCl)-mediated depolarization (**Fig. 1c-1f** and **Extended Data Fig. 1c**). Both first and second phases of insulin secretion were reduced, with the first phase showing a more significant reduction (**Extended Data Fig. 1d-1e**). Inhibition of insulin secretion by low glucose and epinephrine was similar between ND and T2D islets, as was insulin content (**Fig. 1g** and **Extended Data Fig. 1f**); as such, normalization of response by islet insulin content showed similar reductions in stimulated insulin secretion but also showed reduced basal insulin secretion (**Extended Data Fig. 1g-1l**). Together, these data suggest that short-duration T2D islets ex vivo maintain insulin production and storage but have defects at multiple steps of the insulin secretory pathway, including those distal to glucose metabolism, which persist after islet isolation from the in vivo environment.

In contrast to insulin secretion, neither basal nor stimulated glucagon secretion was different in T2D islets when normalized by islet volume (**Fig. 1h-1h** and **Extended Data Fig. 1m**), and both ND and T2D islets showed glucose-mediated suppression of glucagon secretion (**Extended Data Fig. 1n**). Glucagon content was similar between islets from ND and T2D individuals and normalization by glucagon content showed similar secretion dynamics (**Fig. 1L** and **Extended Data Fig. 1o-1t**). While there is substantial evidence of dysregulated glucagon secretion in T2D^28, 29^, these data suggest that either α cell dysfunction is not present in the early stages of T2D or defects are present in vivo but not maintained after islet isolation.

Correlation of donor attributes to functional metrics highlighted a significant negative correlation between donor HbA1c and stimulated insulin secretion (*r*<-0.40, p<0.05; **Fig. 1m**). To test whether the systemic environment contributed to β cell dysfunction in T2D islets, we transplanted T2D or ND islets from a subset of donors into normoglycemic, non-insulin resistant immunodeficient NOD-*scid*-*IL2rγ^null^* (NSG) mice (**Fig. 1n**). After six weeks in this environment, T2D islets secreted less human insulin than ND islets, especially after stimulation with glucose/arginine (**Fig. 1o**, average per donor and **Extended Data Fig. 1u**, individual mice), consistent with ex vivo findings of impaired stimulated insulin secretion. In sum, these experiments highlight that β cell dysfunction in early T2D persists in a normoglycemic, non-insulin resistant environment and suggest that intrinsic β cell dysregulation and/or cellular and molecular alterations within the islet microenvironment are key features driving reduced insulin secretion.

### Broad transcriptional dysregulation revealed through integrated transcriptome analysis of islets and purified α and β cells

To assess both the β and α cell-specific transcriptional landscapes as well as global islet dysregulation in the short-duration T2D cohort, we purified β and α cells by fluorescence-activated cell sorting (FACS) using well-characterized cell surface antibodies and hand-picked isolated islets for RNA-sequencing (**Fig. 1p** and **Extended Data Fig. 1v**). Studying sorted β and α cells together with whole islets, which has not been done in prior studies, allowed detailed appreciation of both cell type-specific and islet-wide transcriptional changes in T2D. As collection of these rare tissues spanned more than 3.5 years, we used a latent variable analysis to discern biological variation from technical variation and then examined the datasets by both differential gene expression (**Fig. 1p** and **Extended Data Fig. 1w, 2a-2f**) and gene network analyses. Differential expression analysis yielded 352, 248, and 564 differentially expressed genes in β cells, α cells, and whole islets, respectively (**Extended Data Fig. 2g-2i**), highlighted by genes involved in stimulated insulin secretion in β cells (*G6PC2, GLP1R*) and changes in non-endocrine components in islets (*CXCL8*, *ADAMTS4*). Numerous metabolic and mitochondrial, exocytosis, ion transport and protein secretion pathways were enriched in T2D β cells (**Extended Data Fig. 2j**), while α cell gene changes were in amino acid and steroid signaling pathways and regulation of blood vessel morphology (**Extended Data Fig. 2k**). In T2D islets, cytokine signaling and immune terms were enriched, as were pathways related to ER processing and unfolded proteins (**Extended Data Fig. 2l**). These were less prominent in isolated α or β cells (**Extended Data Fig. 2j-2k**). Despite diverse differentially expressed genes across sample types (**Fig. 1q**), there was considerable overlap at the level of biological pathways in which these genes are involved – among the most enriched across samples were hormone secretion, lipid metabolism, and cilia organization (**Fig. 1r**). In sum, analysis of differential gene expression of sorted β and α cells and whole islets emphasizes common dysregulated pathways among sample types as well as cell-specific transcriptomic changes.

### Short-duration T2D donors do not show significant changes in endocrine cell mass

To understand the context in which these functional and transcriptomic changes occur, we comprehensively evaluated the islet architecture in pancreatic tissue from T2D donors. High-throughput traditional immunohistochemistry (IHC) was applied across pancreas head, body, and tail regions for the entire donor cohort, and in parallel, a subset of samples was analyzed with a 28-marker panel using co-detection by indexing (CODEX) (**Fig. 2a**). This multiplexed technique for fluorescence-based imaging of large tissue sections without tissue destruction provided simultaneous visualization of multiple tissue compartments as well as spatially resolved cellular phenotypes defined by combined expression/exclusion of multiple markers (**Extended Data Fig. 3a-3b**).

**Figure 2.**
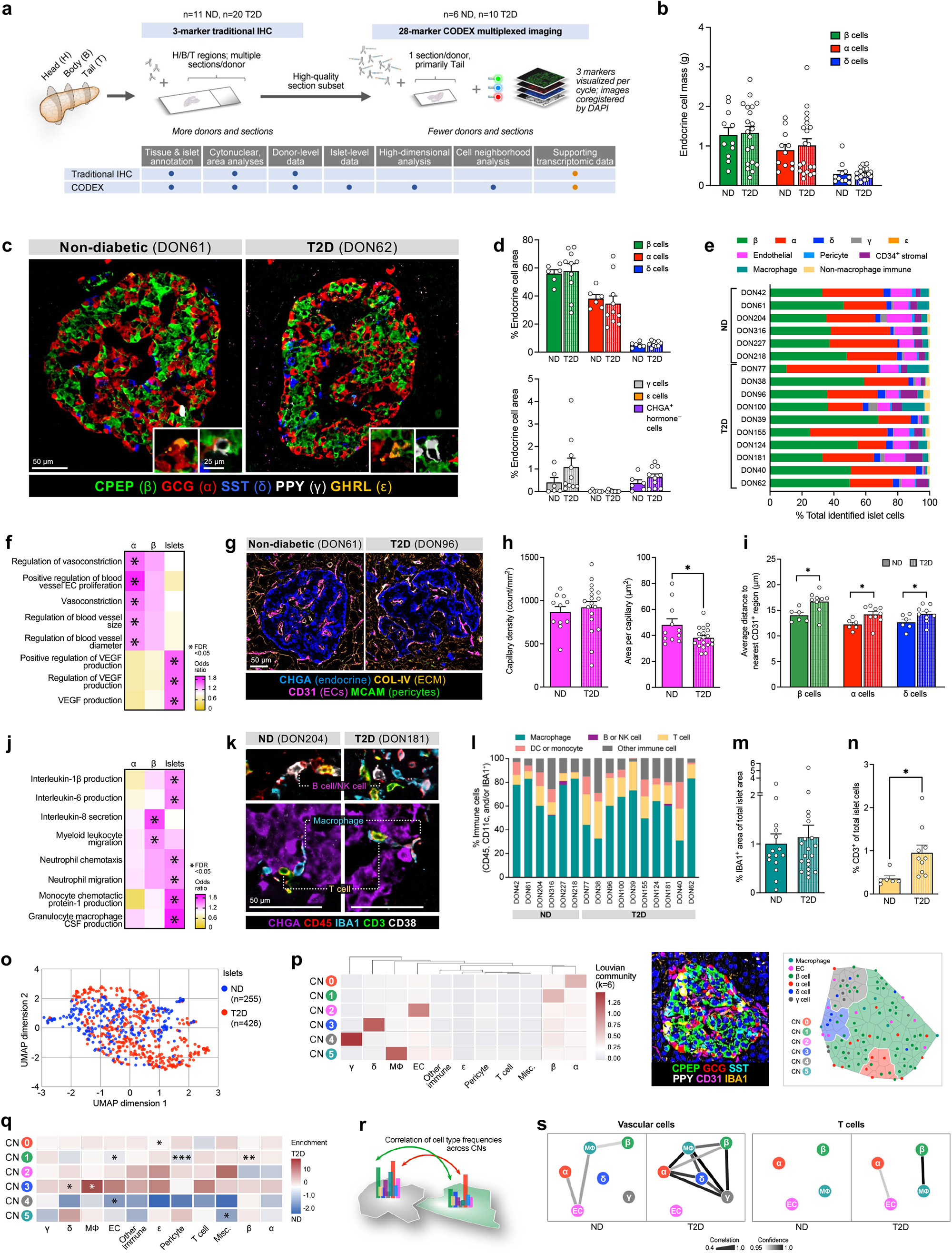
Integrated tissue analysis reveals no change to endocrine cell mass or number, but alteration in intraislet capillaries, T cells, and cellular neighborhoods in short-duration T2D cohort. **(a)** Schematic illustrating parallel analysis by traditional and multiplexed immunohistochemistry (IHC). **(b)** Mass of β, α, and δ cells in ND and T2D donors. **(c)** Representative images of islets from co-detection by indexing (CODEX) imaging; insets show γ and ε cells. **(d)** Cross-sectional area of endocrine cell types. **(e)** Relative proportions of islet endocrine, vascular, stromal, and immune cells. **(f)** Enrichment of vascular-related ontology terms in T2D transcriptome. **(g)** Representative images of islet capillaries, pericytes, and extracellular matrix (ECM). **(h)** Islet capillary density and area per capillary. **(i)** Spatial analysis of endocrine cells and islet capillaries. **(j)** Enrichment of immune-related ontology terms in T2D transcriptome. **(k-l)** Islet immune cell phenotypes and composition. **(m-n)** Islet macrophage **(m)** and T cell **(n)** abundance. **(o)** High-dimensional component analysis of islet cell composition per islet (n=255 ND, n=426 T2D). **(p-s)** Cellular neighborhood assignment **(p)** and corresponding cell composition and correlation changes in T2D vs. ND islets **(q-s)**. Traditional IHC data: panels **b, h, m**; CODEX data: panels **c-e, g, i, k-l, n-s**. Symbols in bar graphs represent donors; * p<0.05 (two-tailed t-test, ND vs. T2D). RNA data: panels **f, j**; * FDR<0.05.

Because changes in endocrine cell number or ratio could explain the reduced insulin secretion in T2D islets, we first evaluated β, α, and δ cell populations. Multiple analyses across pancreas head, body, and tail, including evaluation of islet cell area and islet cell count within entire cross-sections, revealed that β and α cell mass in short-duration T2D were similar to controls (**Fig. 2b** and **Extended Data Fig. 3c-3h**), supporting the similar insulin content in the two groups of islets. We additionally assessed cell death and found apoptotic and/or necrotic cells to be exceedingly rare in both ND and T2D islets (data not shown). Donor-to-donor variability in β and α cell ratio was notable underscoring the challenge in working with heterogeneous human tissues. CODEX permitted simultaneous assessment of rarer γ and ε cell populations as well as identification of cells positive for chromogranin A (CHGA) but negative for all hormones, previously suggested to define “dedifferentiated” β cells^30, 31^. These cells were rare but present in both ND and T2D at similar proportions (**Fig. 2c-2d** and **Extended Data Fig. 3i**). Evidence of amyloid deposits, the abnormal buildup of β cell-produced islet amyloid polypeptide (IAPP) that manifests in T2D, was detectable in 75% of donors in this cohort but with variable prevalence and did not correlate to endocrine cell abundance or area (**Extended Data Fig. 4a-4b**). Thus, tissue analysis suggests that changes in endocrine cell numbers, including β cell mass, are not a substantial component of short-duration T2D. Instead, these data point to reduction in β cell function as the predominant feature of this disease stage.

### Reduced capillary size, increased T cell populations, and altered cellular neighborhoods highlight alterations in T2D islet microenvironment

Adequate islet vascularization and blood flow are critical for sensing and delivery of hormones to systemic circulation, so we next investigated islet capillary endothelial cells (ECs), the most abundant non-endocrine islet cell population (**Fig. 2e** and **Extended Data Fig. 3j**). Pathway analysis from RNA-seq highlighted enrichment in T2D samples for processes controlling blood vessel size, particularly in α cells, as well as regulation of growth factors critical to islet capillary maintenance (**Fig. 2f** and **Extended Data Fig. 4c**). Morphometric analysis demonstrated that capillary size, but not density, was reduced in T2D islets (**Fig. 2h-2i**), resulting in a greater distance of endocrine cells to the nearest capillary in T2D islets (**Fig. 2i**). Interestingly, α and δ cells were closer to capillaries than β cells in both ND and T2D islets (**Extended Data Fig. 4d**), aligning with α cells expressing more angiogenic ligands and receptors than β cells (**Extended Data Fig. 4e**). Phenotypic markers CD34, a cell adhesion molecule that is prevalent in progenitor capillary ECs^32^, and HLA-DR, a major histocompatibility class II (MHCII) receptor, were unchanged in T2D ECs (**Extended Data Fig. 4f**).

In addition to vasculature-related processes, transcriptional profiling also revealed enrichment in T2D β cells and islets for cytokine signaling and immune cell recruitment pathways (**Fig. 2j** and **Extended Data Fig. 4g**). Macrophages, the largest population of intraislet immune cells, did not differ between ND and T2D based on either abundance or phenotypic classification by proinflammatory (HLA-DR^+^) or anti-inflammatory (CD163 and/or CD206^+^) markers (**Fig. 2k-2m** and **Extended Data Fig. 4h**). T cells were rarer in the islet than macrophages but elevated in T2D islets across CD4^+^ (helper), CD8^+^ (cytotoxic), and CD4^−^ CD8^−^ (double negative) phenotypes (**Fig. 2n** and **Extended Data Fig. 4i**). HLA-DR^+^ T cells, previously observed in T2D islets^33^, were not increased, though they were more abundant in a subset of T2D donors (**Extended Data Fig. 4j**). High dimensional data analysis using Uniform Manifold Approximation and Projection (UMAP) of all identified cell types within individually annotated islets revealed a high degree of overlap between islets from ND and T2D donors, emphasizing that although there are subtle differences, the overall islet composition is similar (**Fig. 2o**).

Because analyses of islet composition did not consider the spatial organization of islet cells, we next applied two neighborhood analyses in parallel to annotated islet regions in an effort to identify differential cell architecture. A community detection algorithm tailored to islet cell frequencies, termed CF-IDF, categorized six different cellular neighborhoods (CNs), clusters of cells with distinct cell type compositions that were defined by the most enriched cell type (CN0-CN5; **Fig. 2p**). A modified *k*-means clustering algorithm previously developed for CODEX data corroborated CN classifications (**Extended Data Fig. 4k**), and both approaches found similar CN distribution between ND and T2D islets (**Extended Data Fig. 4l**). ECs and pericytes were depleted in β CNs (CN1) of T2D islets (**Fig. 2q** and **Extended Data Fig. 4m**), consistent with our findings of decreased proximity between β cells and ECs in T2D. In contrast, T2D β CNs had higher β cell enrichment than ND (**Fig. 2q**). We also asked whether cell type frequencies correlated between CNs, i.e., if there was evidence for connectivity between spatially distinct regions (**Fig. 2r**). Vascular cell frequencies were correlated between more CNs in T2D compared to ND islets, while T cell frequencies were specifically correlated between EC and α CNs as well as β and macrophage CNs in T2D (**Fig. 2s** and **Extended Data Fig. 4n**), congruent with findings by islet RNA-seq that EC-specific and immune signals were upregulated in T2D. Together, these results demonstrate modest disruptions of islet organization by vascular and immune cells in early-stage T2D.

### Co-expression network analyses identified gene modules related to donor and islet traits and revealed disrupted metabolism and cilia homeostasis in T2D

To understand the key gene networks that were contributing to β cell dysfunction in short-duration T2D, we performed weighted gene co-expression network analysis (WGCNA) on α cell, β cell, and islet samples (**Extended Data Fig. 6**). This approach created modules (“eigengenes”) of up to 2,000 genes each, labeled by sample type and numbered consecutively (β cells, modules β00-β48; α cells, α00-α54; islets, i00-i67). Collapsing the expression patterns across >14,000 genes into a smaller number of modules reduced gene-level multiple testing burden and enabled association of transcriptomic profiles with sample features including donor traits, islet functional parameters from the same donors defined by dynamic islet perifusion, and enrichment of open chromatin peaks to overlap GWAS variants (β cells: **Fig. 3a-3e**; α cells: **Extended Data Fig. 6a-6e**; islets: **Extended Data Fig. 6f-6i**). Modules with significant correlations were then queried, based on their member genes, for ontology terms to determine biological processes related to significant associations.

**Figure 3.**
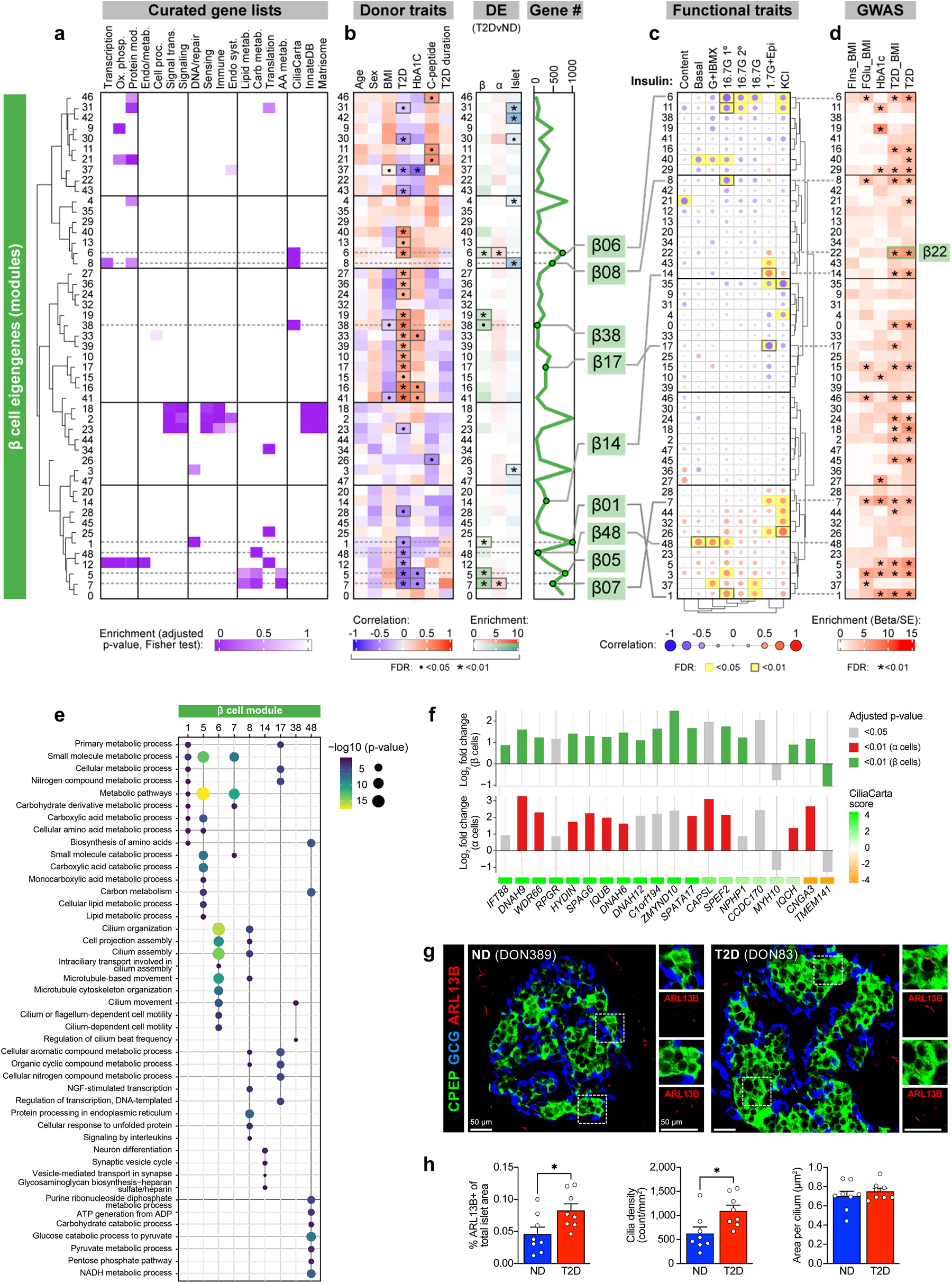
Weighted Gene Co-expression Network Analysis (WGCNA) distinguishes β cell gene modules associated with donor and islet traits as well as those enriched in GWAS loci and identifies disruption in cilia processes as a conserved feature across sample types. **(a)** Relative enrichment of β cell module eigengenes for curated gene lists, based on genes present in each module. **(b)** Module correlation to donor characteristics, enrichment of differentially expressed (DE) genes, and total number of genes per module. • p<0.05; * p<0.01. Modules of interest highlighted (green). **(c)** Module correlation to β cell function described in Fig. 1; significant associations highlighted (yellow). G+IBMX, 16.7 mM glucose with 100 μM isobutylmethylxanthine; 16.7G, 16.7 mM glucose; 16.7G 1°, first phase; 16.7G 2°, second phase; 1.7G+Epi, 1.7 mM glucose and 1 μM epinephrine; KCl, 20 mM potassium chloride. **(d)** Module enrichment for GWAS traits. FIns, fasting insulin; FGlu, fasting glucose. * FDR<0.01. **(e)** Enrichment of select gene ontology terms in β cell modules with notable correlations and/or enrichment. **(f)** Cilia-related genes with fold change ≥ |1.5| in both α and β cells in T2D. **(g-h)** Visualization by immunohistochemistry of cilia (ARL13B; red) and quantification of abundance, density, and size in ND and T2D tissue. * p<0.05 (two-tailed t-test).

Several β cell modules were significantly (FDR < 5%) associated with whole-body glucose homeostasis (HbA1c), and some of these, such as β05 and β07, were also significantly enriched for genes differentially expressed in T2D β cells (**Fig. 3b**). Both β05 and β07 contained genes related to carbohydrate, lipid, and amino acid metabolism (**Fig. 3a** and **3e**), with β07 significantly correlating with KCl-mediated insulin secretion (*r*=0.49, p=0.027; **Fig. 3c**). Modules significantly positively correlated with glucose-stimulated insulin secretion (GSIS) included β01, β03, and β48, all enriched for metabolism-related processes, while β06 and β08, both enriched for cilium movement and motility, were significantly negatively correlated to GSIS (**Fig. 3c** and **3e**). Importantly, aligning functional correlations with enrichment for GWAS loci (**Fig. 3d**) enabled identification of modules that are more likely to be disease-causing (e.g., β01, β03) as opposed to those without GWAS enrichment (e.g., β48) that may instead represent disease-induced transcriptional changes. Thus, this approach allows linking of transcriptional profiles to islet physiological parameters and facilitates prioritization of signatures based on T2D genetic risk.

Though α cell modules showed weaker correlations to donor and functional traits than did β cells, several modules were significantly enriched for cilia-related genes and α08 was also enriched for α cell genes differentially expressed in T2D α cells (**Extended Data Fig. 6a-6b**). Both α08 and α16 significantly inversely correlated with epinephrine-mediated glucagon secretion and were closely related across functional parameters (**Extended Data Fig. 6c**), with α08 showing significant enrichment for T2D GWAS variants (**Extended Data Fig. 6d**). In addition to genes enriched for cilia processes, α08 also included genes related to cytokine signaling and immune response (**Extended Data Fig. 6e**). Similarly, several islet modules showed notable enrichment for immune- and matrisome-related genes (**Extended Data Fig. 6f**); of these, i25 correlated positively with T2D status and inversely with basal insulin secretion and GSIS, while i26 correlated inversely with KCl-mediated insulin secretion (**Extended Data Fig. 6g-6h**). Genes in both modules corresponded to cell-cell communication, including response to stimulus (i26) and leukocyte activation and migration (i25) (**Extended Data Fig. 6i**). Overall, these patterns suggest that β cell function may be influenced by α and other non-endocrine cells residing within the islet.

Interestingly, cilia-related processes not only defined key functionally correlated modules in every sample type, but they were also some of the most enriched pathways across all samples based on differential gene expression (**Extended Data Fig. 6j**). Further β06, β08, and α08 were enriched for T2D and related trait GWAS loci, suggesting a potential casual role (**Fig. 3d** and **Extended Data Fig. 6d**). We compared fold change of validated cilia-related genes^34^ and determined that the majority were expressed at higher levels in T2D compared to ND for both β and α cells (**Fig. 3f**). To investigate whether these changes translated to cellular alterations, we stained tissue sections from the same donors with cilia marker ARL13B (**Fig. 3g**). Total cilia area within the islet was greater in T2D tissue, attributable to a higher cilia density with unchanged cilia size (**Fig. 3h**), consistent with elevations in cilia transcripts. Thus, integration of functional, transcriptional, genetic, and tissue-based analyses highlights cilia-related processes as playing a key role in early T2D.

### β cell hub gene RFX6 is reduced in T2D and controls glucose-stimulated insulin secretion

The network approach of WGCNA enables identification of “hub” genes that are highly connected, i.e., whose expression highly correlates with many other genes, both within and across modules, making it a powerful analysis to understand central transcriptional regulators that may be driving β cell dysfunction in short-duration T2D (**Fig. 4a**). Of the highly connected β cell genes, *RFX6* stood out as a key islet-enriched transcription factor that has been linked to both monogenic and polygenic forms of diabetes^15, 35, 36^ and thus is in prime position to exert disproportionate influence on the β cell transcriptional state. *RFX6* was more highly connected than other islet-enriched transcription factors specifically in β cells (**Fig. 4a-4b** and **Extended Data Fig. 7a-7d**) and was one of the most reduced islet-enriched transcription factors at the transcript level in T2D β cells (**Fig. 4c**). Importantly, *RFX6* is a member of module β01, which had the strongest positive association with high glucose-stimulated insulin secretion and was among the most significantly enriched for both GWAS variants and RFX binding motifs (**Fig. 3c-3d** and **4d**). Immunohistochemistry analysis revealed a reduction in number of β cells expressing RFX6 in T2D (**Fig. 4e-4f**). Together, these data support RFX6 as a critical hub gene in β cells that may contribute to the functional deficits observed in short-duration T2D.

**Figure 4.**
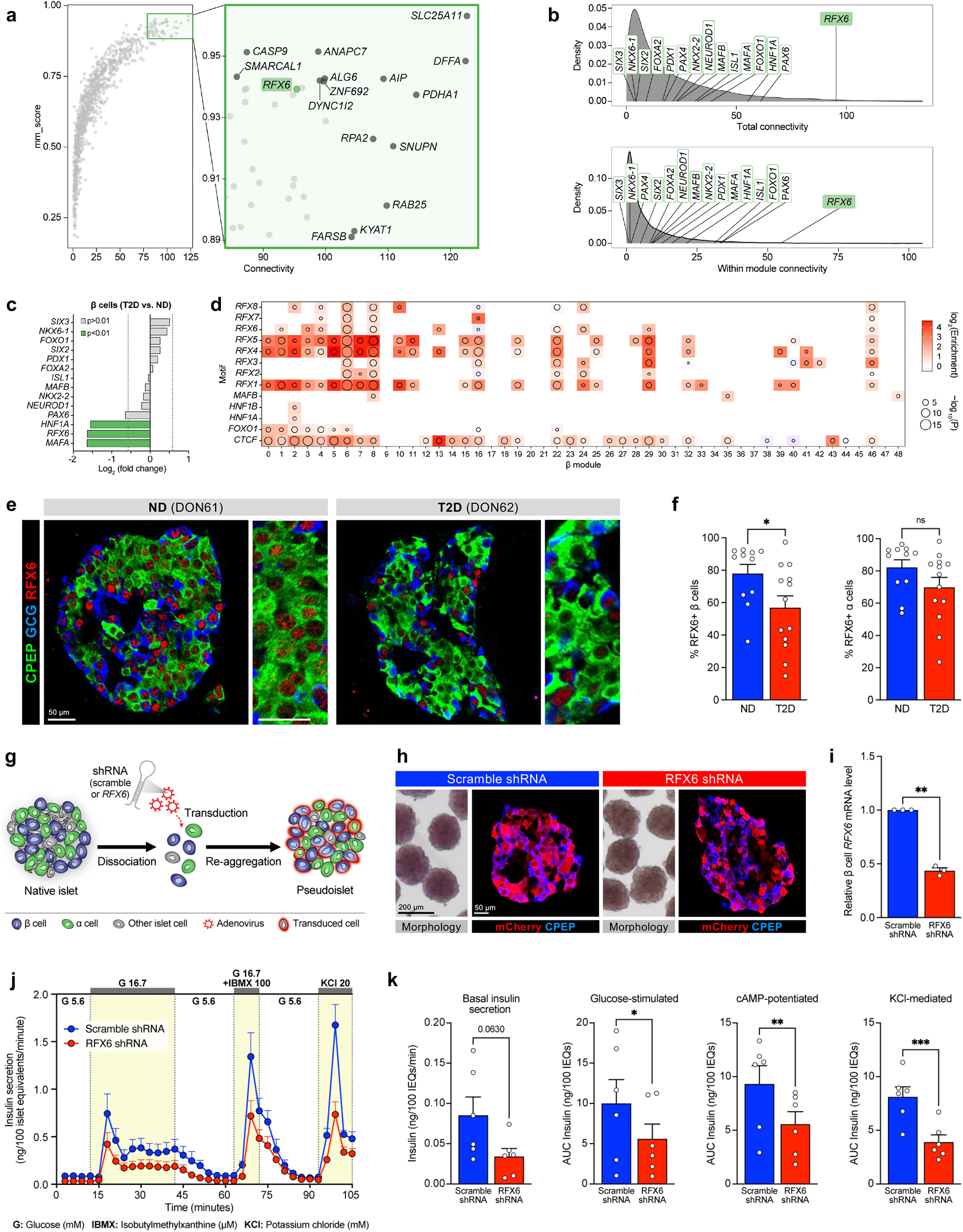
RFX6, a central regulator of transcript changes in short-duration T2D, is reduced in T2D β cells and controls stimulated insulin secretion. **(a-b)** Overall connectivity (**a**) and cross- and within-module connectivity (**b**) of individual genes based on β cell WGCNA. Select genes with high connectivity scores (**a**) and select transcription factors (**b**) are labeled. **(c)** RNA fold change in T2D β cells of transcription factors highlighted in panel **b**. Vertical lines denote fold change = |1.5|. **(d)** Enrichment of transcription factor motifs in β cell modules. **(e-f)** Expression of RFX6 in β and α cells of ND and T2D donors. **(g)** Schematic of adenoviral shRNA delivery and formation of pseudoislets. **(h)** Morphology and immunofluorescent staining of transduced pseudoislets. **(i)** Relative *RFX6* mRNA expression in β cells treated with scramble or *RFX6* shRNA. **(j)** Pseudoislet insulin secretion assessed by perifusion; n=6 donors per group. **(k)** Area under the curve (AUC) for secretory response to each of the stimuli shown in panel **j**. Panels **f**, **i**, **k**: * p<0.05, ** p<0.01, *** p<0.001 (two-tailed t-test).

To determine the role of RFX6 in adult human β cell function in an islet-like context, we used shRNA knockdown in a primary human pseudoislet system that allows for functional and transcriptomic assessment (**Fig. 4g**). Scramble shRNA (‘control’) and *RFX6* shRNA (‘sh*RFX6*’) pseudoislets exhibited similar size and morphology, and preferential β cell transduction resulted in β cell *RFX6* knockdown that did not change β or α cell proportion (**Fig. 4h-4i** and **Extended Data Fig. 7e-7g**), suggesting that acute (6-day) reduction of *RFX6* expression does not lead to β cell loss. Following RFX6 knockdown, dynamic insulin secretion in the presence of three secretagogues (high glucose, high glucose + IBMX, and KCl) was significantly blunted, similar to that seen in T2D islets (**Fig. 5j-5k**). Normalization to insulin content, which was greater in sh*RFX6* pseudoislets, made this secretory response even more prominent (**Extended Data Fig. 7h-7j**). In sum, not only is RFX6 decreased in T2D β cells, but the results of targeted knockdown are consistent with the *RFX6*-containing module β01 association with glucose-stimulated insulin secretion (**Fig. 3d**) and strongly implicate RFX6 as a major regulator of stimulated insulin secretion.

### RFX6 knockdown alters the β cell chromatin and transcriptional landscape and downregulates secretory vesicle components

To determine the molecular mechanism by which RFX6 knockdown impacted insulin secretion, sh*RFX6* and control pseudoislets (n=7 matched donors) were multiplexed using a blocked study design and processed for single nucleus multiome profiling (**Fig. 5a**). Single nucleus (sn)RNA and snATAC reads were collected and filtered to yield 15,825 (RNA) and 5,706 (ATAC) high-quality nuclei for downstream analysis (**Extended Data Fig. 8a**). Islet cell types were resolved by clustering (**Fig. 5b-5c** and **Extended Data Fig. 8b**) where we found representation of all major cell types across all donors (**Extended Data Fig. 8c**) and equal distribution between sh*RFX6* and control constructs (**Fig. 5d**). Consistent with the previously observed preferential adenoviral targeting of β relative to α cells, fluorescent reporter expression was much higher in β cell nuclei than in α cell nuclei (**Fig. 5e**).

**Figure 5.**
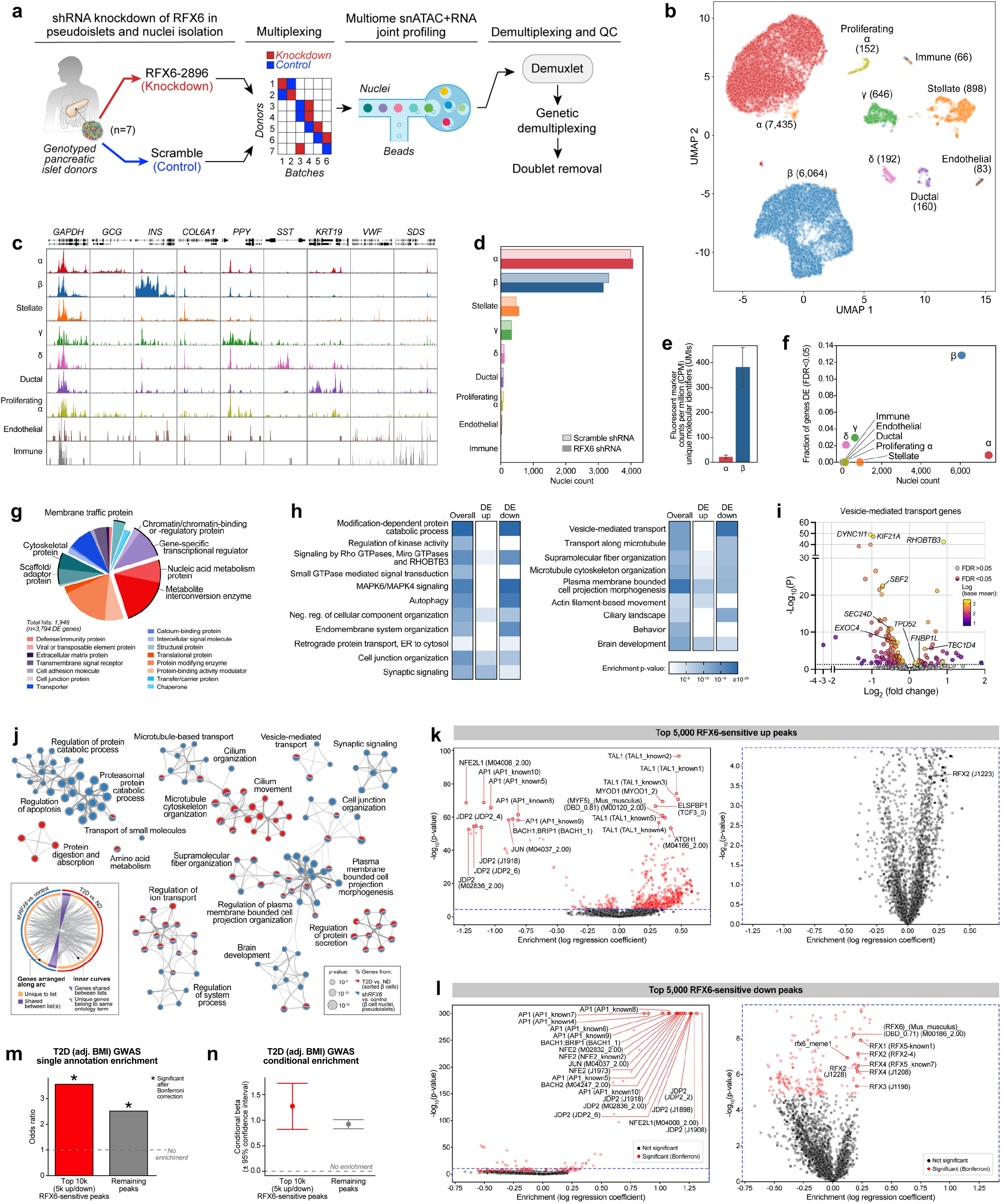
RFX6 controls glucose-stimulated insulin secretion in human β cells through chromatin modifications and vesicle trafficking pathways. **(a)** Schematic depicting randomized study design to mitigate batch effects in single nuclear (sn) RNA- and ATAC-sequencing of scramble shRNA (control) and RFX6 shRNA (sh*RFX6*) pseudoislets. **(b)** Cell type assignment by clustering on RNA. **(c)**. Pseudobulk ATAC signal at marker genes. **(d)** Post-QC nuclei counts from control and sh*RFX6* pseudoislets. **(e)** Abundance of fluorescent marker gene expression (mCherry/mKate2) in α and β cell nuclei. **(f)** Proportion of differentially expressed (DE) genes per cell type. **(g)** Classification of protein-coding DE genes in sh*RFX6* β cells by PANTHER. **(h)** Pathway enrichment for DE genes (FDR<0.01); second two columns separate genes up- or downregulated in sh*RFX6*. **(i)** DE genes in Reactome pathway R-HSA-5653656. **(j)** Overlap of 1,000 most significant DE genes in sh*RFX6* vs. control β cell nuclei (blue) and T2D vs. ND sorted β cells (red), analyzed by Metascape. Circos plot illustrates overlap at the level of genes (purple) or ontology terms (grey). Network displays a subset of enriched terms, where edges denote term similarity and node colors represent contribution of each gene list. **(k-l)** Motif enrichment for top 5,000 RFX6-sensitive up-**(k)** and downregulated **(l)** ATAC peaks in sh*RFX6* β cell nuclei. Right panels show enlarged views of plots on left. **(m-n)** Odds ratio of T2D GWAS enrichment **(m)** and model estimate from conditional analysis **(n)** of RFX6-sensitive peaks.

Supporting the role of RFX6 as a major β cell regulator, 13% of total detected genes were differentially expressed in β cell nuclei compared with <3% in other cell types (**Fig. 5f**). Nuclear *RFX6* was not among those reduced, consistent with shRNA silencing occurring in the cytoplasm. Differentially expressed genes included those encoding cytoskeletal and scaffold/adaptor proteins (11% of those classified), membrane traffic proteins (4%), and gene-specific transcriptional regulator or chromatin/chromatin-binding or -regulatory proteins (13%) (**Fig. 5g**). Upregulated genes were enriched for actin filament-based movement and synaptic signaling, while downregulated genes were enriched for membrane trafficking, autophagy, and ciliary pathways (**Fig. 5h-5i**). To investigate overlap in differentially expressed genes between sh*RFX6* β cell nuclei and sorted T2D β cells, we compared the top 1,000 most significantly differential genes in each group and observed common pathway enrichment related to microtubule cytoskeleton organization, ion transport, and regulation of protein secretion (**Fig. 5j**). Also of note, sh*RFX6* β cell nuclei differentially expressed genes were overrepresented in WGCNA module β22 (**Extended Data Fig. 8d**) that was enriched for T2D GWAS variants and RFX binding motifs. Genes in this module corresponded to cellular membrane and vesicle components, mirroring pathways dysregulated in sh*RFX6* β cell nuclei (**Extended Data Fig. 8e**) and further implicating exocytosis as a target of RFX6-mediated dysfunction in T2D β cells.

We next sought to identify the landscape of chromatin alterations in sh*RFX6* β cells and observed global changes compared to matched controls (**Extended Data Fig. 8f-8g**). We took n=2,000-10,000 peaks with smallest p-values in either direction (‘top RFX6-sensitive peaks’) for use in downstream analyses. These peaks were significantly enriched for motifs corresponding to the known chromatin modifier activator protein 1 (AP1), as well as RFX6 and related family member motifs (**Fig. 5k-5l** and **Extended Data Fig. 8h-8i**). CCCTC-binding factor (CTCF) and RFX motif footprint signatures like those previously observed in bulk islet ATAC data^15^ confirmed the high quality of the snATAC data (**Extended Data Fig. 8j**). Further, top RFX6-sensitive peaks were significantly enriched to occur near differentially expressed genes (**Extended Data Fig. 8k**), indicating concordance between the snATAC and snRNA modalities. We and others have shown that β cell ATAC peaks are enriched for T2D GWAS variants^5, 6^, and indeed, top RFX6-sensitive peaks were also significantly enriched to overlap with these variants (**Fig. 5m** and **Extended Data Fig. 8l**). Importantly, enrichment remained significant after conditional analysis controlled for remaining (not RFX6-sensitive) peaks (**Fig. 5n** and **Extended Data Fig. 8m**), which emphasizes the importance of β cell *RFX6*-sensitive peaks in the genetic predisposition to T2D. Overall, these results show that knockdown of *RFX6* in β cells results in widespread transcriptional and chromatin changes that are associated with downregulated vesicle transport and coordinated disruption of regulatory elements that overlap T2D GWAS variants, consistent with the role of *RFX6* as a master regulator of β cell identity.

## DISCUSSION

The pancreatic β cell, a major focus in diabetes, exists within the multicellular pancreatic islet mini-organ, where interactions between various cell types are increasingly recognized. In T2D, like in other chronic, complex, multi-organ diseases, teasing apart the causes, correlates, and consequences of cellular and tissue dysfunction is challenging due to limited availability of primary tissue, constraints of sample processing at different disease stages, and in many cases, removal of cells from their native environment. To address these challenges and identify early disease-driving events, we applied a comprehensive, multimodal, integrated approach to isolated islets and pancreatic tissue from a unique cohort of short-duration T2D and control donors that included analyses of islet physiology, transcriptome, and pancreas tissue cellular architecture. Furthermore, we integrated donor and islet functional traits with gene network analysis and GWAS to understand central transcriptional regulators driving β cell dysfunction in short-duration T2D. Co-registration of multimodal data and clinical information yielded several important findings (**Extended Data Fig. 9a**): (1) impaired β cell function, a hallmark of early-stage T2D, persisted ex vivo and in nondiabetic environments; in contrast, α cell function was not changed; (2) islet endocrine composition was unchanged though there were modest alterations to the islet microenvironment in endothelial and immune cells; (3) transcriptional network analysis proportioned genetic risk into gene modules with specific functional properties, and (4) *RFX6* emerged as a highly connected hub transcription factor that was reduced in T2D β cells and associated with reduced glucose-stimulated insulin secretion. We validated a critical role for RFX6 by performing dynamic functional analyses and integrated snRNA and snATAC-seq on primary human pseudoislets with knockdown of *RFX6* in β cells. Reduction of RFX6 led to reduced insulin secretion defined by transcriptional dysregulation of vesicle trafficking, exocytosis, and ion transport pathways that was mediated by chromatin architectural changes overlapping with T2D GWAS variants (**Extended Data Fig. 9b**). Thus, our integrated, multimodal studies identify β cell dysfunction that results from cell-intrinsic defects, including an RFX6-mediated, T2D GWAS-enriched transcriptional network, as a key event in early T2D pathogenesis.

### Dysfunction of β cells, and not β cell loss, is primary defect in early-stage T2D

This study demonstrates β cell functional defects ex vivo – which persist in culture and following transplantation into a normoglycemic environment – but no change to insulin content or β cell mass. The relative contributions of impaired β cell function and/or reduced β cell mass have long been debated in T2D^37–39^. Though postmortem studies suggest mild β cell loss^33, 40–43^, most studies mixed short- and long-term disease duration together and noted that defects were more severe with longer duration and/or insulin treatment. Recent studies of metabolically profiled donors suggested that β cell loss is not prominent in early T2D^26, 44^. By integrating studies of both pancreatic tissue and isolated islets from the same donors, our data indicate that β cell loss is not a major component in disease pathogenesis at early-stage T2D. Further, the continued dysfunction of islets in a transplant setting also underscores the persistence of initial β cell defect. In sum, this study illustrates that β cell dysfunction occurs early in T2D and that prevention and/or rapid intervention may be critical to preserve β cell function.

### Changes to islet microenvironment emphasize additional disease processes that may become more prominent in later disease stages

Our transcriptional analyses in isolated islets identified altered vascular and immune signaling as features in sorted α and β cells as well as in whole islets. Although isolated islets do not provide a physiologic context, particularly for endothelial cells without their connection to systemic circulation, similar transcriptional changes were found in laser capture microdissected T2D islets^26^. Further, our comprehensive tissue analyses of the same donors allowed in situ characterization of non-endocrine islet cell abundance, phenotype, and localization. We demonstrated that T2D islets had subtle reductions in islet capillary size, increased intraislet T cells, and altered communication between cellular neighborhoods, but overall the microenvironment was largely similar to ND islets. While most donors showed some evidence of amyloid deposits as a unique feature of the T2D islet microenvironment, only a minority of islets demonstrated detectable amyloid at this stage of disease. Together, these observations are unlikely to explain the degree of β cell dysfunction in this cohort but, given that they are present without any associated changes in endocrine cell composition, may represent early consequences of β cell dysfunction or may act to exacerbate initial β cell-intrinsic defects.

Indeed, inflammatory signals and other trophic factors have been shown to influence β cell function, especially in the presence of amyloid, and may become a more prominent feature of the disease at later stages^19, 45–47^. Further study is needed to determine whether changes to the microenvironment are truly an independent disease process or whether there is bidirectional signaling between dysfunctional β cells, α cells, and/or other islet cell types.

### Integrated co-expression network analysis reveals gene modules of genetic risk in T2D

The transcriptomic profiles of sorted α and β cells in addition to islets provided new insight into cell-specific contributions to T2D pathogenesis. Co-expression network analysis and association with GWAS variants and physiological parameters, similar to a recent approach^48^, allowed us to prioritize processes with physiological relevance that were more likely to be disease-causing rather than disease-induced. For instance, both β01 (metabolism-enriched) and β06 (cilia-enriched) modules are associated with T2D GWAS variants, indicating that regulatory circuitry related to metabolism and cilia function may have causative roles in development of T2D. Notably, insulin secretion was positively correlated to β01, whose genes were decreased in T2D β cells, but negatively correlated to β06, whose genes were increased in T2D β cells. These results suggest that β01 genes enhance insulin secretion while β06 genes decrease it, thus one expects that T2D risk alleles likely decrease β01 gene expression and activate β06 genes, both of which would negatively influence β cell function. Future work directly testing key candidate genes from this dataset, analogous to the studies of RFX6 described here, will be important to validate these processes and how they contribute to T2D pathogenesis.

Genetic risk for complex metabolic diseases such as T2D results from the combined influence of many small-effect variants, with at-risk individuals likely having multiple parallel processes affected. This concept has been described as a “palette” model^49^, and our work aids in deciphering components of the palette by proportioning genetic risk into cell-specific functional modules derived from transcriptome signatures across early stages of disease. Thus, this opens the opportunity to assess downstream consequences of an individual’s innate genetic risk by identifying specific molecular and functional processes that would be most affected and hopefully allowing for precise targeting of those to achieve personalized medicine in diabetes.

#### RFX6 plays a central role in dysregulation of β cell function early in T2D

By identifying an RFX6 regulatory network that strongly correlates with insulin secretion and T2D genetic risk, this study provides new insight into previous work which has linked RFX6 to both monogenic and polygenic forms of diabetes^15, 35, 36^. Our results suggest that RFX6 exerts a disproportionate transcriptional influence on β cell state and that its dysregulation is a key molecular event in early T2D pathogenesis. We pursued this finding by directly testing the role of RFX6 in pseudoislets and demonstrated a clear function for RFX6 in governing stimulated insulin secretion in primary human β cells. Previous studies with direct perturbation of RFX6 in adult β cells, performed in cell lines and mouse models, highlighted downstream effects on Ca^2+^ and KATP channels^50, 51^. Our work confirms defective ion transport processes but identifies vesicle trafficking and exocytosis pathways as major drivers of defective insulin secretion in primary human β cells with impaired release likely responsible for the buildup of insulin content. Additionally, we show that these transcriptional changes are mediated by changes in β cell chromatin regions significantly overlapping with T2D GWAS loci, emphasizing the central role of RFX6 in mediating genetic risk to functional defects that define early T2D. Further, cilia-related genes were also significantly dysregulated following RFX6 reduction, in line with evidence that the RFX family of transcription factors control ciliogenesis^52, 53^. Given their role in environment sensing, cell-to-cell communication, and signal transduction, cilia represent a potential link between β cell-intrinsic, RFX6-mediated dysregulation and changes within the islet microenvironment seen in early T2D and warrant future study.

This work raises important questions about what factors or events initially dysregulate RFX6 to start this cascade. Given the coordinating role RFX6 plays in islet cell development^35^, it may be that early defects driven by RFX6 dysfunction only become apparent after superimposed environmental, nutritional, and/or age-related stressors. Alternatively, the strong enrichment of T2D GWAS variants in β01 (the *RFX6*-containing module) and position of RFX6 as a hub gene may point to cumulative genetic effects compounding over time in an irreversible cascade that disrupts β cell homeostasis. Thus, precisely what underlies the initial dysregulation of RFX6, and whether it can be targeted to prevent or reverse early molecular defects in the β cell, should be an active area of investigation.

## METHODS

### Human subjects

Pancreata from nondiabetic (ND) (n=19) and T2D (n=20) donors were obtained through partnerships with the International Institute for Advancement of Medicine (IIAM), National Disease Research Interchange (NDRI), and local organ procurement organizations. Pancreata were processed in Pittsburgh by Dr. Rita Bottino for both islet isolation and histological analysis as previously described^1–3^. Additional ND human islet preparations (n=27) were obtained through partnerships with the Integrated Islet Distribution Program (IIDP) and Alberta Diabetes Institute (ADI) Isletcore and served as assay-specific controls or were used for pseudoislet studies. Donor information is detailed in Extended Data Table 1. Deidentified medical histories provided both information for T2D staging as well as clinical characteristics to correlate with generated data. The Vanderbilt University Institutional Review Board declared that studies on de-identified human pancreatic specimens do not qualify as human subject research.

Some human islets used in this research study were provided by the ADI IsletCore at the University of Alberta in Edmonton (http://www.bcell.org/adi-isletcore.html) with the assistance of the Human Organ Procurement and Exchange (HOPE) program, Trillium Gift of Life Network (TGLN), and other Canadian organ procurement organizations. Islet isolation was approved by the Human Research Ethics Board at the University of Alberta (Pro00013094). All donors’ families gave informed consent for the use of pancreatic tissue in research. This study also used data from the Organ Procurement and Transplantation Network (OPTN) that was in part compiled from the Data Hub accessible to IIDP-affiliated investigators through IIDP portal https://iidp.coh.org/secure/isletavail). The OPTN data system includes data on all donors, wait-listed candidates, and transplant recipients in the US, submitted by the members of the OPTN. The Health Resources and Services Administration (HRSA), U.S. Department of Health and Human Services provides oversight to the activities of the OPTN contractor. The data reported here have been supplied by UNOS as the contractor for the Organ Procurement and Transplantation Network (OPTN). The interpretation and reporting of these data are the responsibility of the authors and in no way should be seen as an official policy of or interpretation by the OPTN or the U.S. Government.

### Pancreas procurement and processing

Pancreata from ND and T2D donors (see **Extended Data Table 1** for donor information) were received within 18 hours from cross clamp and maintained in cold preservation solution on ice until processing, as described previously^3^. Pancreas was then cleaned from connective tissue and fat, measured, and weighed. Prior to islet isolation, multiple cross-sectional slices of pancreas with 2-3 mm thickness were obtained from the head, body and distal tail, further divided into quadrants, and processed into paraformaldehyde-fixed cryosections as described previously^3^. Islet isolation was performed via ductal collagenase infusion and purification by density gradient as described previously^1, 3^, then shipped to Vanderbilt for further analysis following shipping protocols developed by the IIDP. Islets were cultured in CMRL 1066 media (5.5 mM glucose, 10% FBS, 1% Pen/Strep, and 2 mM L-glutamine) in 5% CO2 at 37°C for 24– 48 hours prior to reported studies^3–5^. Pseudoislets were cultured in Vanderbilt Pseudoislet media^6^. Limitations of tissue availability and processing dictated that not all assays could be performed on each donor.

### Assessment of native pancreatic islet and pseudoislet function by macroperifusion

Function of islets from ND and T2D donors and pseudoislets were studied in a dynamic cell perifusion system at a perifusate flow rate of 1 mL/min^6, 7^. The effluent was collected at 3-minute intervals using an automatic fraction collector, then islets were retrieved and lysed with acid-ethanol solution to extract. Insulin and/or glucagon concentrations in each perifusion fraction, as well as total hormone content, were measured by radioimmunoassay (RIA) (human insulin, RI-13K, Millipore; glucagon, GL-32K, Millipore), enzyme-linked immunosorbent assay (ELISA) (Human insulin, 10-1132-01, Mercodia; glucagon, 10-1281-01, Mercodia), or Homogeneous Time Resolved Fluorescence (HTRF) assay (glucagon, 62CGLPEH, Cisbio). Area under the curve (AUC) above baseline hormone release was calculated with the trapezoidal method in GraphPad Prism 8.0-9.3 as previously described^6^.

### Human islet transplantation

Immunodeficient NOD.Cg-*Prkdc^scid^Il2rg^tm1Wjl^*/Sz (NSG)^8^ 10- to 12-week old male mice were maintained by Vanderbilt Division of Animal Care in group housing in sterile containers within a pathogen-free barrier facility housed with a 12 hour light/12 hour dark cycle and access to free water and standard rodent chow. All animal procedures were approved by the Vanderbilt Institutional Animal Care and Use Committees. Between 1000-2000 islet equivalents per mouse (n=4-8 mice per islet preparation) were transplanted beneath the kidney capsule. After 6 weeks, mice were fasted for 6 hours and then injected with glucose + arginine (2g/kg body weight) intraperitoneally as previously described^3–5, 9^. Blood samples were obtained before (0’) and after (15’) injection and human-specific insulin was analyzed by ELISA (Alpco, 80-ISNHU-E01.1) or radioimmunoassay (Millipore, RI-13K).

### Purification of α and β cells by FACS

Human islets from ND and T2D donors were dispersed and sorted for collection of RNA from α and β cells as described previously^3, 10^. Briefly, 0.025% trypsin was used to disperse islet cells by manual pipetting and subsequently quenched with RPMI containing 10% FBS. Cells were washed in the same medium and counted on a hemocytometer, then transferred to FACS buffer (2 mM EDTA, 2% FBS, 1X PBS). Indirect antibody labeling was completed via two sequential incubation periods at 4C, with one wash in the FACS buffer following each incubation. Primary and secondary antibodies, listed in **Extended Data Table 2**, have been characterized previously and used to isolate high-quality RNA^3, 10–12^. Appropriate single color compensation controls were run alongside samples. For sorting of β cells for use in pseudoislets, quenching step post-dispersion was performed with 100% FBS at 1/3 volume trypsin. Cells then underwent an additional filtration step using a 40 μl strainer prior to staining. For all preparations, propidium iodide (0.05 ug/100,000 cells; BD Biosciences, San Jose, CA) was added to samples prior to sorting for non-viable cell exclusion. Flow analysis was performed using an LSRFortessa cell analyzer (BD Biosciences, San Jose, CA), and a FACSAria III cell sorter (BD Biosciences, San Jose, CA) was used for FACS. Cells for RNA were collected into FACS buffer, washed once in 1X PBS, and stored in RNA lysis buffer for RNA extraction. Cells for pseudoislets were washed once in 1X PBS, resuspended in Vanderbilt pseudoislet media, and processed as described in *Pseudoislet* section below. Analysis of flow cytometry data was completed using FlowJo 10.1.5 (Tree Star, Ashland, OR).

### Traditional and multiplexed immunohistochemical imaging and analysis

#### Traditional Immunohistochemistry

Multiple sections from pancreatic head, body, and tail regions of 20 T2D and 11 age-matched ND donors were lightly paraformaldehyde (PFA)-fixed and prepared for immunohistochemistry and stained as described previously^3, 10, 13^. Primary and secondary antibodies and their dilutions are listed in **Extended Data Table 2**. Amyloid was visualized using a 2-minute incubation in Thioflavin S (0.5% w/v; #T-1892, Sigma, St. Louis, MO) followed by a brief wash in 70% ethanol as described previously^5, 9, 14^. Images were acquired at 20X with 2X digital zoom using a FV3000 confocal laser scanning microscope (Olympus) or a ScanScope FL (Aperio) and processed using cytonuclear algorithms (HighPlex FL v3.2.1) or tissue classifiers via HALO software (Indica Labs) or morphometric measurement via Metamorph software v7.10 (Molecular Devices, LLC). Analyses were run on the entire tissue section or manually annotated islets as indicated in figure legends. Endocrine cell mass was quantified by using pancreas weight and the ratio of hormone positive cells as identified by cytonuclear algorithm within the entire pancreatic section from multiple blocks representing the head, body, and tail regions. To obtain islet capillary measurements, caveolin-1 channel was isolated and color thresholding was used on a per-image basis to gather object data using the Integrated Morphometry Analysis (IMA) function (Metamorph). The following analysis metrics represent mean ± standard error: endocrine cells (**Fig. 2b, Extended Data Fig. 3f-3h**) 16,151 ± 1,715 islet cells/donor and 570,508 ± 51,866 total cells/donor; endocrine cell area (**Extended Data Fig. 3c-3d**) 2.34 ± 0.24 mm^2^/donor; capillary morphology (**Fig. 2h**) 48 ± 4 islets/donor; macrophage area (**Fig. 2m**) 0.64 ± 0.07 mm^2^/donor; amyloid (**Extended Data Fig. 4a**) 108 ± 19 islets/donor; cilia (**Fig. 3h**) 0.32 ± 0.05 mm^2^/donor; RFX6 (**Fig. 4f**) 1,863 ± 362 cells/donor; pseudoislets (**Extended Data Fig. 7g**) 2,797 ± 508 cells/sample.

#### CODEX multiplexed imaging

Antibodies were purchased preconjugated from Akoya Biosciences or sourced from other vendors and conjugated in-house using the CODEX Conjugation Kit (Akoya Biosciences) or by Leinco Technologies, Inc. (St. Louis, MO, USA) (**Extended Data Table 3**). 10-μm lightly fixed^3^ pancreas sections were mounted onto 22x22 mm glass coverslips (Electron Microscopy Sciences) coated in 0.1% Poly-L-lysine (Sigma) and stained with the CODEX Staining Kit (Akoya Biosciences) in uncoated 6-well tissue culture plates (VWR) per manufacturer instructions. Fluorescent oligonucleotide-conjugated reporters were combined with Nuclear Stain and CODEX Assay Reagent (Akoya Biosciences) in light-protected 96-well plates sealed with foil (Akoya Biosciences) and automated image acquisition and fluidics exchange were performed using the Akoya CODEX instrument and CODEX Instrument Manager (CIM) v1.29 driver software (Akoya Biosciences) integrated with a BZ-X800 epifluorescent microscope (Keyence). Tissue was hydrated in 1X CODEX buffer (10X CODEX Buffer diluted in Milli-Q water) and hybridization/stripping of the fluorescent oligonucleotides was performed using dimethyl sulfoxide (Sigma). After loading of coverslip into stage insert, tissue was visualized with Nuclear Stain diluted 1:1000 in PBS and imaging area was set by center point and tile number using BZ-X800 viewing software (Keyence). All images were acquired using a CFI plan Apo I 20x/0.75 objective (Nikon) with 30% tile overlap and 5 z-planes (1.5 μm/z).

#### Processing and annotation of CODEX images

A total of 16 tissue regions were captured from 6 ND and 10 T2D donors (mean 50 mm^2^ tissue/donor). Image alignment, stitching, background subtraction, and deconvolution were performed using the CODEX Processor v1.7.0.6 (Akoya Biosciences; see https://help.codex.bio/codex/processor/technical-notes for details). Individual channel images (TIFF files) were imported into HALO software v3.1 (Indica Labs) for all analyses as described below. Tissue and islet areas were annotated by hand to exclude out-of-focus regions and poor tissue quality. Islets (estimated diameter ≥50 μm; mean 42 islets/donor) were annotated based on DAPI and CHGA channels. Cell segmentation and cell type annotations were performed using the HALO HighPlex FL v3.2.1 module with consistent cytonuclear parameters (nuclear contrast threshold 0.456, maximum cytoplasm radius 0.48). Due to marker intensity variability among samples, thresholds were manually set for each marker and donor. Unless otherwise noted, cells were counted positive for a given marker if minimum intensity was reached in 50% of cytoplasm area (see **Extended Data Fig. 3a-3b** for complete list of markers, abbreviations, and cell types). For cells with more variable morphology, positivity was also counted for nuclear area (30%: ARG1, CD11c, CD14, CD163, CD206, CD31, CD34, CD45, HLA-DR, IBA1, KRT, MCAM). Proliferating cells were counted only if minimum 60% of nuclear area met Ki67 intensity threshold. Vascular structures (CD31) were also measured by random forest classification algorithm (HALO Tissue Classifier module). The following analysis metrics represent mean ± standard error: endocrine cell area (**Fig. 2d**) 0.88 ± 0.10 mm^2^/donor; islet cell composition (**Fig. 2e**, **Extended Data Fig. 3j**) 7,322 ± 852 cells/donor; immune cells (**Fig. 2l, 2n**) 309 ± 43 cells/donor; endothelial cell phenotypes (**Extended Data Fig. 4f**) 460 ± 92 cells/donor; macrophage phenotypes (**Extended Data Fig. 4h**) 191 ± 29 cells/donor; T cell phenotypes (**Extended Data Fig. 4i-4j**) 40 ± 17 cells/donor.

#### High-dimensional, spatial, and neighborhood analyses

The R implementation of the UMAP algorithm (https://CRAN.R-project.org/package=umap) was used for dimensionality reduction. Cell marker percentages obtained through HALO were standardized across islets (n=255 ND islets and 426 T2D islets; mean 172 cells/islet), and default parameters were used for UMAP reduction (**Fig. 2o**) except for nearest neighbors (80) and minimum distance (0.05). For spatial analyses, CD31 area classifications were converted to an annotation layer. A nearest neighbor algorithm (HALO Spatial Analysis module) was applied to obtain average distance of endocrine cells (n=4,830 ± 692 cells/donor) to islet capillaries (CD31^+^ region) (**Fig. 2i**, **Extended Data Fig. 4d**).

For cell neighborhood (CN) analysis, two methods were applied in parallel to CODEX data from annotated islets. In the community detection method, termed *Dynamic CF-IDF* (**Fig. 2p-2q, 2s**), a weighted undirected heterogeneous graph for each islet was constructed based on the cell types and normalized distance between cells. A greedy-based graph community detection method^15^ was applied to segment the graph into a set of cell communities, then cell communities were stratified into 6 CNs (n=5,582 total CNs with median 11 cells/CN). Cell type enrichment was determined by a new proposed scoring function *CF-IDF*, which is a modification of the widely used text sequence analysis method term frequency (TF)–inverse document frequency (IDF) scoring^16^. Our cell frequency (CF)-inverse dataset frequency (IDF) score emphasizes the cell type that is not only prevailing, but also uniquely representative in a group of target islets. Therefore, it will deemphasize the most dominant cell types (e.g., α and β) throughout all the islets while paying more attention to the relative enrichment of less abundant cell types (e.g., vascular and immune cells) in the local regions. The downstream analysis not only introduces insightful results on T2D feature analysis but also shows a robust performance across different resolution levels.

The second CN analysis method, a *k*-means approach (**Extended Data Fig. 4k-4n**), built on a previously published algorithm used to identify CNs in the tumor microenvironment^17^. For each cell, we first found its 10 nearest neighbors in the islet and assigned the *i-th* nearest neighbor which was an α cell, β cell, macrophage, EC cell, or γ cell, a score *cos(iπ/20)*. Then we calculated the total score for each cell type, applied L1 normalization to the scores, and standardized them across all cells. The resulting representations of cells were finally used for *k*-means clustering to form 5 CNs (n=5,021 total CNs with median 5 cells/CN).

### Transcriptional analysis of α and β cells and islets from ND and T2D donors

#### RNA isolation and bulk RNA-sequencing

RNA was extracted from sorted α and β cells (see above, *Purification of α and β cells by* FACS) or from pelleted whole islets using the Invitrogen RNAqueous-Micro Total RNA Isolation kit (Thermo Fisher #AM1931). TURBO DNA-free (Ambion) was used to treat any trace DNA contamination. RNA was quantified by Qubit Fluorometer 2.0 and RNA integrity was confirmed (RIN >7) by 2100 Bioanalyzer (Agilent). Amplified cDNA libraries were constructed using SMART-seq v4 Ultra low Input RNA-kit (Takara) and sequencing was performed on an NovaSeq platform (Illumina) using paired-end reads (100 bp) and 25 million reads per sample.

We processed the raw RNA-seq reads using FastQC (v0.11.8) for broad quality assessment. Briefly, we examined the following parameters: (1) base quality score distribution, (2) sequence quality score distribution, (3) average base content per read, (4) GC distribution in thereads, (5) PCR amplification issue, (6) overrepresented sequences, (7) adapter content. Based on the quality report of fastq files, we trimmed sequence reads using fastq-mcf (v1.05) and cutadapt (v2.5) to only retain high quality sequence for further analysis. The paired-end reads were aligned to the GRCh37/hg19 human reference with GENCODE v19 gene annotation using STAR splice-aware aligner (v2.5.4b; --outSAMUnmapped Within KeepPairs)^18^.

We counted fragments mapping to features type in GENCODE v19 gene annotation using featureCounts from Subread package^19^. The gene list was pruned to contain only protein-coding genes mapping to autosome and chrX, resulting in a total of 20,260 genes. We assessed libraries using comprehensive quality metrics generated by QoRTs^20^ as well as computed derived metrics. Briefly, on the top of QoRTs reported metrics, we computed (1) 5’-3’ gene coverage bias (as the ratio of coverage values at the 90%-ile and 10%-ile of the coverage distribution), (2) Kolmogorov-Smirov test statistic between cumulative gene diversity of each library relative to median distribution of all libraries within each cell type and standardized to a mean of 0 and standard deviation of 1 to yield a z-score, (3) number of reads mapped mapped to *Xist* and *SRY* genes, (4) average number of reads mapped to chrM, and (5) transcript integrity number (TIN)^21^ for each library. The labeled sex of donors was matched against the gene expression quantified for sex genes to rule out any sample swaps or mislabeling. We also computed principal components for TPM normalized count matrix for each cell type in order to detect potential outliers.

#### Differential gene expression analysis

We performed differential gene expression analysis between T2D and ND samples for each cell type individually using DESeq2^22^. In order to minimize potential effects of known and unknown confounding factors, we included known covariates in the DESeq2 model as well accounted for unknown covariates using RUVseq latent variable approach^23^. Briefly, we used the following multi-step process: (1) We first removed genes from the raw count matrix which had less than 10 reads in fewer than 25% of the samples for that cell type. (2) We then ran a first-pass differential expression analysis using DESeq2 with Age, Sex, BMI, and Batch as known covariates. The output result was filtered for genes that were non-significant i.e., not differentially expressed between T2D and ND samples and had p-value > 0.5. These genes were used as “control” or “empirical” genes for RUVSeq::RUVg function to estimate latent variables accounting for variation in the data not attributed to disease status. (3) The latent variables estimated from the RUVseq run were then used as additional covariates (on the top of Age, Sex, BMI, and Batch where applicable) for the second run of DESeq2. We selected the number of latent variables to provide the most reasonable separation between T2D and ND samples and minimal deviation from mean in the relative log expression plots. The output results from DESeq2 were filtered for 1% FDR to generate the final list of genes differentially expressed between T2D and ND for each cell type. We performed functional enrichment analysis using RNA-enrich^24^ and retained terms with an FDR threshold of 5%. Terms were condensed using the RelSim function in REVIGO^25^ with similarity parameter set to 0.5 and visualized in semantic space using an.xgmml file imported into Cytoscape software^26^ v3.8.2.

Combined analysis of differentially expressed genes (fold change ≥1.5 or ≤-1.5; p<0.01) was performed using Metascape^27^ v3.5. Metascape’s heuristic algorithm samples the 20 top-score clusters, selects up to the 10 best scoring terms (lowest p-values) within each cluster, and connects terms pairs with Kappa similarity above 0.3. The resulting network was exported as a .cys file and visualized using Cytoscape, with the most representative term name in each cluster selected manually.

#### Gene network analysis

We adopted Weighted Gene Co-expression Network Analysis (WGCNA)^28^ approach to create networks from the gene expression data. Briefly, we first filtered genes following the same rule established in Differential Gene Expression analysis where we only kept genes that had at least 10 reads in at least 25% of the samples for each cell type. We then processed raw counts using the varianceStabilizedTransformation function in DESeq2 package and used removeBatchEffect from the limma R package^29^ to adjust for effects of age, sex, and BMI while protecting for disease status in the design matrix. The normalized and batch corrected count matrix was then used as input to blockwiseModules to create a “signed hybrid” network with “bicor” as the correlation function. The power (k) parameter was selected such that the scale free topology fit reached at least 80% fit. To examine cell type modules associated with quantitative traits of interest, we utilized a linear regression-based framework. We (1) inverse normalized the raw quantitative trait, (2) adjusted for Age, Sex, and BMI by linear regression, and (3) computed the spearman rank correlation between residuals and eigengene of all modules. Within each network, we also computed the module membership score and network connectivity for each gene. Estimated enrichment of curated gene lists^30–32^ (**Extended Data Table 4**) was calculated using Fisher’s exact test. Functional enrichment of genes in each module was performed using gprofiler2^33^, and the results were visualized as a dotplot.

#### Integration of network analysis with chromatin accessibility

We integrated chromatin accessibility information with gene network analysis using sci-ATAC-seq data for α and β cells derived from our previously published study^34^. For each module within each cell type, we selected (a) accessible sites that were present within a specified distance of the transcription start site (TSS) of the genes within that module, and (b) the distal chromatin peaks that were linked to the peaks within this set based on the Cicero peak interaction results from the same study. This set of TSS proximal and distal peaks for all of the genes within each module and for each cell type were then used for downstream enrichment analyses.

For variant enrichment analysis in the module linked peaks, we collected the latest published summary statistics for selected traits^35, 36^. Using a threshold of +-10kb to define our gene TSS boundary for linking peaks with modules, we created a set of accessible sites for each module. The union of peaks across all modules was used as a “bulk” positive enrichment control. We then tested the enrichment of trait-associated variants from multiple GWAS across module peaks using GARFIELD^37^ and used a p-value threshold of 5e-08 as input parameter for selecting trait-associated variants.

Next, we considered whether specific Transcription Factor Binding Motifs (TFBMs) are enriched to occur in certain modules. To test this, we defined module linked peaks for each module as described before but using a threshold of +-1kb from gene TSS. For each peak within a module, we then identified the peak summit and extended the summit by 50 bp in each direction. Using genomic sequence in this region as our “test sequence”, we used Analysis of Motif Enrichment (AME, v5.3.2) tool from MEME-Suite^38^ (using default parameters) to identify enriched TFBMs represented in cisBP v.2.0^39^. The control set of sequence was generated using --shuffle--parameter in AME which generates a control sequence by shuffling the test sequence but preserving the 2-mer frequency. The enrichment score was computed as scaled log2 transformed (TP+1)/(FP+1) for each TFBM.

### Pseudoislet formation and assessment of RFX6 knockdown

Pseudoislets were formed as previously described^6^. Briefly, nondiabetic human islets were handpicked to purity and then dispersed with 0.025% HyClone trypsin (Thermo Scientific) for 7 minutes at room temperature before counting with an automated Countess II cell counter or manually by hemacytometer. Dispersed human islets or purified β cells (see above, *Purification of α and β cells by FACS*) were incubated in adenovirus at a multiplicity of infection of 500 for 2 hours in Vanderbilt pseudoislet media before being spun and washed. Adenovirus containing U6 driven scramble or RFX6 targeted shRNA as well as CMV driven mCherry or mKate2 red fluorescent tag were prepared, amplified and purified by Welgen, Inc (Worcester, MA). Cells were then resuspended in appropriate volume of Vanderbilt pseudoislet media to allow for seeding into wells at 2000 cells per 200 µL each well of CellCarrier Spheroid Ultra-low attachment microplates (PerkinElmer). Pseudoislets were allowed to reaggregate for 6 days before being harvested and studied.

To assess knockdown, RNA was extracted from pseudoislets containing only β cells using an RNAqueous RNA isolation kit (Ambion). cDNA synthesis and quantitative reverse transcriptase PCR were performed as previously described^2^; briefly, cDNA was synthesized using a High-Capacity cDNA Reverse Transcription Kit (Applied Biosystems #4368814) according to the manufacturer’s instructions. Quantitative PCR (qPCR) was performed using TaqMan probes for ACTB (Hs99999903_m1) as endogenous control and RFX6 (Hs00941591_m1). Relative changes in mRNA expression were calculated by the comparative ΔCt method.

### Multiome single nuclear RNA/ATAC-sequencing

#### Nuclear isolation

Pseudoislet samples treated with RFX6 shRNA or scramble RNA were pooled together using a randomized study design, so the targeting and scramble conditions were not confounded by batch (**Fig. 5a**). To accomplish this, samples were allocated into six groups (batches) of n=490-494 pseudoislets for nuclei isolation. A customized protocol was developed based on recommendations by 10x Genomics (https://www.10xgenomics.com/resources/demonstrated-protocols/) which included optimization steps described below. Briefly, the samples were suspended in 1X PBS and pelleted at 2000 x g for 3 minutes at 4°C. The pellet was resuspended in lysis buffer (10mM Tris-HCl 7.4 pH, 10mM NaCl, 3mM MgCl2, 0.1% Tween-20, 0.1% NP40, 0.01% Digitonin, 1% BSA, 1mM DTT, and 2U/µl RNase Inhibitor) and rocked in an Eppendorf thermomixer C (EP #5382000015) at 300 x g for 5 minutes at 4°C. Keeping the samples on ice as much as possible, tubes were then transferred to a prechilled 2 mL glass dounce homogenizer and homogenized with 15 strokes of tight pestle B before being transferred to a 1.5 mL tube and centrifuged at 500 x g for 5 minutes at 4°C. The resulting pellet was then resuspended in 1 mL of wash buffer (10mM Tris-HCL 7.4 pH, 10mM NaCl, 3mM MgCl2, 1% BSA, 0.1% Tween-20, 1Mm DTT, and 2U/µl RNase Inhibitor) and centrifuged at 100 x g for 1 minute at 4°C. The supernatant was collected, filtered through a pre-wetted 30 µm filter, and centrifuged at 500 x g for 5 minutes at 4°C. Nuclei were resuspended in 300 µl of wash buffer, then 300 µl of sucrose cushion (0.88M sucrose, 1mM DTT, 1mM RNAse Inhibitor, and 10% wash buffer) was added to the bottom of the tube and the resulting layered solution was centrifuged at 1000 x g for 10 minutes at 4°C. Both layers of supernatant were removed, and pellet was resuspended in 1 mL of wash buffer and centrifuged at 500 x g for 5 minutes at 4°C. Nuclei were then resuspended in 30 µl of nuclei resuspension buffer before counting and quality assessment. The desired concentration of nuclei was achieved by resuspending the appropriate number of nuclei in 1X diluted nuclei buffer for joint (on the same nucleus) snATAC-seq and snRNA-seq multiome profiling. Nuclei were processed by the University of Michigan Advanced Genomics Core using the 10x Genomics Chromium platform at 20K nuclei per well.

#### Multiome sample genotyping and imputation

Samples were genotyped with the Infinium Multi-Ethnic Global-8 v1.0 kit using 50 ng/uL DNA samples in two batches. Probes were mapped to Build 37. We merged the .ped files for the two batches along with samples from other projects that were genotyped on the same chip (resulting in a combined 68 samples). We removed variants with multi mapping probes and updated the variant rsIDs using Illumina support files Multi-EthnicGlobal_D1_MappingComment.txt and Multi-EthnicGlobal_D1.annotated.txt (downloaded from https://support.illumina.com/downloads/infinium-multi-ethnic-global-8-v1-support-files.html). We performed pre-imputation QC using the HRC-1000G-check-bim.pl script (version 4.2.9) obtained from the Mark McCarthy lab website (https://www.well.ox.ac.uk/∼wrayner/tools/) to check for strand, alleles, position, Ref/Alt assignments and update the same based on the 1000G reference (https://www.well.ox.ac.uk/∼wrayner/tools/1000GP_Phase3_combined.legend.gz).

We did not conduct allele frequency checks at this step (i.e. used the --noexclude flag) since we had 68 samples from mixed ancestries. These filters resulted in 958,427 variants. We performed pre-phasing and imputation using the Michigan Imputation Server^40^. The standard pipeline (https://imputationserver.readthedocs.io/en/latest/pipeline/) included pre-phasing using Eagle2^41^ and genotype dosage imputation using Minimac4 (https://github.com/statgen/Minimac4) and the 1000g phase 3 v5 (build GRCh37/hg19) reference panel^42^. Post-imputation, we selected biallelic variants with estimated imputation accuracy (r∧2) > 0.3, variants not significantly deviating from Hardy Weinberg Equilibrium (P>1e-6), MAF in 1000G European individuals > 0.05 and minor allele count (MAC) > 1 in our 12 samples, resulting in 6,665,607 variants.

#### Data processing (RNA component)

The RNA component of the multiome data was processed using starSOLO (STAR v. 2.7.3a, with GENCODE v19 annotation; options --soloUMIfiltering MultiGeneUMI --soloCBmatchWLtype 1MM_multi_pseudocounts --soloCellFilter None), which outputs the count matrices needed for most of the analyses^18^. Quality control metrics were gathered on a per-nucleus basis using a custom Python script on the corrected gene counts and aligned BAM file.

Following processing with STAR, we constructed a custom count matrix by combining information from the GeneFull and Gene matrices output by STAR. The GeneFull matrix contains per-gene counts based on intronic and exonic reads, while the Gene matrix contains per-gene counts based on exonic reads only. As nuclear RNA may contain introns, the GeneFull matrix should be preferred. However, due to overlapping transcript annotations that render some read gene assignments ambiguous, some genes may receive fewer counts in the GeneFull matrix than in the Gene matrix. The INS gene was an extreme example of this, receiving very low counts in the GeneFull matrix but high counts in the Gene matrix. To salvage counts for such genes, our custom matrix utilized the GeneFull counts for most genes but utilized the Gene counts for the subset of genes that had greater counts in the Gene matrix than in the GeneFull matrix.

#### Data processing (ATAC component)

Adapters were trimmed using cta (https://github.com/ParkerLab/cta). We used a custom Python script, available in the Parker lab Github repository, for barcode correction. Barcodes were corrected in a similar manner as in the 10x Genomics Cell Ranger ATAC v. 1.0 software. In brief, barcodes were checked against the 10x Genomics whitelist. If a barcode was not on the whitelist, then we found all whitelisted barcodes within a hamming distance of two from the bad barcode. For each of these whitelisted barcodes, we calculated the probability that the bad barcode should be assigned to the whitelisted barcode using the Phred scores of the mismatched base(s) and the prior probability of a read coming from the whitelisted barcode (based on the whitelisted barcode’s abundance in the rest of the data). If there was at least a 97.5% probability that the bad barcode was derived from one specific whitelisted barcode, it was corrected to the whitelisted barcode.

Reads were mapped using BWA-MEM^43^ with flags ‘-I 200,200,5000 -M’ (v. 0.7.15-r1140). We used Picard MarkDuplicates (v. 2.25.1; https://broadinstitute.github.io/picard/) to mark duplicates, and filtered to high-quality, non-duplicate autosomal read pairs using SAMtools view^44^ with flags ‘-f 3 -F 4 -F 8 -F 256 -F 1024 -F 2048 -q 30’ (v. 1.10). Quality control metrics were gathered on a per-nucleus basis using ataqv^45^ (v. 1.2.1) on the BAM file with duplicates marked.

#### Selection of quality nuclei (barcodes) for downstream analysis

We performed rigorous QC of all RNA nuclei and only included those deemed as high-quality based on the following four definitions: 1) nUMI > 1000, 2) mitochondrial fraction < 0.2, 3) nuclei where the RNA profile was statistically different from the background/ambient RNA signal, and 4) nuclei identifiable as a singlet and assignable to a sample using genotypes. We considered droplets with UMIs < 10 to be “empty” and therefore representative of the background/ambient RNA profile. Top genes in the ambient RNA included highly expressed genes across prominent islet cell types such as INS, GCG, and SST, along with several mitochondrial genes. We used the testEmptyDrops function from DropletUtils (v 1.6.1)^46^, specifying the ‘lower’ parameter as 10 and selecting droplets with P<0.05 as droplets significantly different from the ambient RNA profile. To identify singlets and assign to samples, we ran Demuxlet^47^ using using the BAM files and the genotype VCF file considering all post-QC variants in gene bodies with minor allele count (MAC) >1. We used the command “demuxlet --sam $bam --tag-group CB --tag-UMI UB --vcf ${vcf} --alpha 0 --alpha 0.5 --field GT”, and selected singlets. To account for ambient RNA contamination while identifying singlets, we also masked the top 1% genes expressed in the ambient RNA and re-ran Demuxlet with the same parameters; nuclei were considered singlets and kept for downstream analysis if they were called as singlets in either Demuxlet run.

We also performed QC of the ATAC component of the multiome data. For ATAC, we required nuclei to have a minimum TSS enrichment (as calculated by ataqv) of 2, minimum filtered read count of 1000 (ataqv ‘HQAA’ metric), and maximum mitochondrial fraction of 0.5. We also ran Demuxlet on the ATAC component (command: demuxlet --sam $bam --tag-group CB --vcf ${vcf} --field GT) and required that a prospective nucleus be called as a singlet. The ATAC component of nuclei in two wells showed low TSS enrichment and all nuclei from these two wells were therefore excluded from analysis.

If the RNA and the ATAC component of a barcode both passed QC and the Demuxlet sample assignment was the same, both modalities were utilized for downstream analysis. If only the RNA component passed QC, only the RNA component was used in downstream analysis. As we performed clustering on the RNA component, we excluded the few (twelve) barcodes that passed ATAC QC and failed RNA QC.

#### Removal of ambient RNA counts from single nucleus gene expression UMI matrices

Prior to clustering and downstream analysis, we used DecontX^48^ (celda v. 1.8.1, in R v. 4.1.1)^49^ to adjust the nucleus x gene expression count matrices for ambient RNA. DecontX was run on a per-batch basis, as the amount of ambient contamination may vary across batches.

Decontaminated counts were generated via the decontX() function, passing barcodes with total UMI count <= 10 to the background argument. Rounded decontaminated counts were used for clustering and all downstream analyses. Nuclei with estimated contamination level > 0.2 were excluded from downstream analysis.

#### Clustering of multiome data

Nuclei were clustered on the RNA component using Seurat^50–52^ (v. 3.9.9.9010, in R v. 3.6.3). After normalizing counts with the NormalizeData function, we identified the top 2000 variable features (FindVariableFeatures function, with selection.method=’vst’) and scaled with the ScaleData function. We identified neighbors using the top 20 PCs and k.param = 20, and called clusters using resolution = 0.1 with n.start = 100. We used the top 20 PCs for generating the UMAP.

This clustering protocol identified 10 clusters. One of the smaller clusters shows expression of both INS and GCG, suggesting it may consist of doublets that were not caught by demuxlet. To verify this was a doublet cluster, we ran a different, genotype-independent, ATAC-based doublet detection method (AMULET; v. 1.0-beta, run with default parameters separately on data from each multiome well)^53^ on the ATAC nuclei that otherwise passed QC. This method tagged ∼40% of the nuclei in the suspected doublet cluster as doublets, while only ∼5% of nuclei in any other cluster were tagged as doublets. We therefore removed the small doublet cluster from the clustering and downstream analysis.

#### Differential gene expression analysis

Differential gene expression was performed within each cluster using DESeq2 (v. 1.28.0)^22^ on pseudobulk counts. UMI counts were summed across nuclei within a donor + construct + cluster. Only donors with paired data (RFX6-2896 and scrambled-mCherry constructs) were used, and the analysis was performed in a paired fashion (DESeq2 model: ∼donor + construct). We used an FDR threshold of 5% for considering genes differentially expressed.

#### Per-cluster processing of ATAC component

All ATAC reads from pass-QC, clustered nuclei were merged within each cluster. To generate per-cluster peaks, these BAM files were converted to single-ended BED format using bedtools bamtobed^54^ before calling ATAC-seq peak summits with MACS2^55^ (flags -g hs --nomodel --shift -37 --extsize 73 -B --keep-dup all --call-summits). We removed summits in blacklist regions, filtered to FDR 0.1% summits, and then generated a peak list from the summits by extending the ATAC-seq peak summits for each cluster +/-150 bps to get 300bp peaks (within each cluster, if two 300bp peaks overlapped the one with the greater MACS2 score was kept). We then removed peaks in blacklist regions. To get the ATAC peak counts used in the ATAC PCA and differential chromatin accessibility analyses, we determined the number of ATAC fragments overlapping each of these peaks in each of the per-cluster, per-donor, per-construct pseudobulk samples.

For visualization of ATAC signal, we generated a normalized bedGraph file using MACS2 on the single-end BED file (macs2 callpeak command, with options --SPMR --nomodel --shift -100 -- extsize 200 -B --broad --keep-dup all) and then converted to bigWig format using the UCSC bedGraphToBigWig^56^. For PCA on the pseudobulk ATAC counts, we first removed any peaks on the mCherry or mKate2 contigs. We then converted peak counts to counts per million and removed the bottom 10% of features with the lowest average CPM across samples. For each peak, we filled any 0s with a value equal to half of the minimum non-zero CPM for that peak across samples. We then log transformed prior to performing the PCA.

#### Differential chromatin accessibility analysis

Differential chromatin accessibility was performed within each cluster using DESeq2 (v. 1.28.0)^22^ on pseudobulk ATAC peak counts. Only donors with paired ATAC data (RFX6-2896 and scrambled-mCherry constructs) were used, and we additionally excluded donor 17277513 due to very low read counts. The DESeq2 analysis was performed in a paired fashion, with model: ∼donor + tss_enrichment + construct. To compute TSS enrichment for each pseudobulk sample, we merged all ATAC nuclei (regardless of cluster) from each donor and computed TSS enrichment with ataqv.

#### Testing for enrichment of peak subsets near differential genes

We used a permutation test to determine whether the most significant peaks (‘top peaks’) from the beta cell differential peak analysis were enriched near beta cell differentially expressed (DE) genes. First, we assigned each peak to the gene with the nearest TSS (if multiple TSS were equally close, we took the TSS with the smallest chromosomal coordinate). We then calculated the fraction of top peaks whose nearest gene was DE. To get the null expectation for this value, we permuted the ‘DE/not DE’ gene labels, such that the same number of genes were always labeled as ‘DE’ but the identity of these DE genes changed in each permutation. While permuting, we split genes into deciles based on the expression of each gene and permuted the labels only within each decile (this controls for the fact that highly expressed genes are more likely to be DE than lowly expressed genes due to statistical power in the DE analysis). We performed 10,000 permutations, in each permutation re-calculating the fraction of top peaks whose nearest gene was DE to build up the null distribution. We then calculated an empirical p-value based on our observed value and the null distribution, adding a pseudocount to avoid a p-value of 0 (p = [1 + # of permutations where the test statistic was greater than or equal to our observed value] / 10,001).

#### Motif scanning for multiome motif enrichment analyses

The motif scans were performed using FIMO (v. 5.0.4) with a background model calculated from the hg19 reference genome^57^ and otherwise default parameters. We used the motifs from Kheradpour and Kellis 2014^58^, excluding “*_disc” motifs; motifs from cisBP v. 2.0^39^; motifs from Jolma et al. 2013^59^; and custom RFX6 motifs generated using mouse Rfx6 ChIP-seq data from Piccand et al. 2014^60^.

The custom RFX6 motifs were generated during a previous project^61^. Sequencing reads from Piccand et al. 1014^60^ were mapped to the mouse mm9 genome^62^ using bwa (v. 0.7.12-r1039) and peaks were called using MACS2 (flags: -t MIN6_Rfx6-HA_IP.bam -c MIN6_Control-HA.bam -B --nomodel -g mm --keep-dup 1 -q 1.00e-4). The MEME (v. 4.11.0)^63^ and DREME (v. 4.9.1)^64^ tools from the MEME suite^65^ were used to discover novel motifs in the resulting peaks. One non-repetitive motif from the MEME tool and two motifs from the DREME tool, bearing similarity to known RFX family motifs, were selected for use in downstream analysis.

#### Motif enrichment in most significant peaks

We used logistic regression to measure enrichment of motifs in subsets of ATAC-seq peaks. We ran one model per peak category and motif. For testing for enrichment in the peaks that had the smallest p-values and leaned towards higher signal in shRFX6 samples, we modeled:

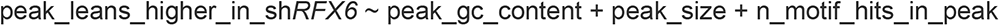

Where ‘peak_leans_higher_in_sh*RFX6’* is 1 if the peak was one of the most significant peaks in the ‘up in RFX6 KD condition’ direction and 0 otherwise; peak_gc_content was the GC content of the sequence within the peak; peak_size was the mean DESeq2-normalized count for the peak across the samples in the DESeq2 analysis; and n_motif_hits_in_peak was the number of motif hits in the peak as determined by the FIMO motif scans. The coefficient of the n_motif_hits_in_peak term was taken as the measure of motif enrichment. For testing for enrichment in the peaks that had the smallest p-values and leaned towards lower signal in sh*RFX6* samples, we used the same model except the outcome variable was ‘peak_leans_lower_in_sh*RFX6*’.

#### Generation of ATAC footprint plots

To generate the ATAC footprint plots, we first separated the motif occurrences into those within the beta cells ATAC peaks and those outside of peaks. For each of these two groups, we computed an aggregate Tn5 cut matrix for the 500 bps on either side of the motifs, using beta cell ATAC reads from each individual donor+construct (using the make_cut_matrix script within the atactk package (https://github.com/ParkerLab/atactk); options -a -r 500). The cut matrices were generated separately for each donor+construct, utilizing only donors with paired ATAC data (RFX6-2896 and scrambled-mCherry constructs) and additionally excluding donor 17277513 due to very low ATAC read counts. To reduce the impact of Tn5 insertion sequence bias, we normalized the Tn5 cut frequency at each position for the motifs in peaks by the corresponding frequencies for the motifs outside of peaks. To adjust for technical differences (e.g., TSS enrichment) between the donors+constructs, we then divided these normalized cut frequencies by the average normalized cut frequency between the -500 and -400 bp positions.

#### GWAS enrichment in most significant peaks

We considered if β cell ATAC-seq peaks that score highly for differential accessibility, as measured by p-value, are specifically enriched to overlap T2D-GWAS variants. We compared the enrichment of T2D (adj. BMI) GWAS variants to overlap top 5000 ATAC-seq differential peaks leaning up and down with the remaining peaks for β cell using GARFIELD^37^. Using a p-value threshold of 1e-05, we also performed a conditional analysis where GARFIELD evaluates if both annotations are conditionally independent of each other in the enrichment model. The coefficients corresponding to each annotation from the conditional enrichment model were shown along with the 95%-CI. To ensure robustness of our results, we repeated the analysis for top 2000 (up and down each) and top 10000 (up and down each) differential peaks.

### Statistical information

Specific statistical tests used for each dataset are described in the figure legends and text where appropriate. Data is represented as mean ± standard error (SEM) unless otherwise noted. A p-value of 0.05 was considered significant except for bulk RNA-seq differential expression where we used a more stringent cutoff of 0.01. Statistical comparisons were performed using GraphPad Prism software 8.0-9.3 or using R.

## ACKNOWLEDGEMENTS

We thank the organ donors and their families for their invaluable donations and the International Institute for Advancement of Medicine (IIAM), Organ Procurement Organizations, National Disease Research Exchange (NDRI), and the Alberta Diabetes Institute IsletCore together with the Human Organ Procurement and Exchange (HOPE) program and Trillium Gift of Life Network (TGLN) for their partnership in studies of human pancreatic tissue for research. We thank Drs. Jing Hughes, Jeong Hun Jo, Seung Kim, Yan Hang, Joana Almaça, Roland Stein, and Rachana Haliyur for their valuable scientific insight regarding experimental design and methods. This study used human pancreatic islets that were provided by the NIDDK-funded Integrated Islet Distribution Program at the City of Hope (DK098085). This work was supported by the Human Islet Research Network (RRID:SCR_014393), the Human Pancreas Analysis Program (RRID:SCR_016202), DK106755, DK123716, DK123743, DK120456, DK104211, DK108120, DK104218, DK112232, DK117960, DK126185, DK117147, EY032442, T32GM007347, F30DK118830, DK20593 (Vanderbilt Diabetes Research and Training Center), The Leona M. and Harry B. Helmsley Charitable Trust, JDRF, Doris Duke Charitable Foundation, and the Department of Veterans Affairs (BX000666). Cell sorting was performed in the Vanderbilt Flow Cytometry Shared Resource (P30 CA68485, DK058404) and whole-slide imaging was performed in the Islet and Pancreas Analysis Core of the Vanderbilt DRTC (DK20593).

## AUTHOR CONTRIBUTIONS

Conceptualization, JTW, DCS, VR, CD, JJW, SCJP, ACP, MB; Software, VR, PO, YT, SF, AV; Formal Analysis, VR, PO, ALH, YT, SF, SS, AV, SA; Investigation, JTW, DCS, CD, ALH, CVR, JJW, YDP, CV, RA, GP, RJ, NJH; Resources, DLG, LDS, RB; Data Curation, JTW, DCS, VR, PO; Writing – Original Draft, JTW, DCS, VR, SCJP, ACP, MB; Writing – Review & Editing, all authors; Visualization, JTW, DCS, VR, PO, SS, AV; Supervision, JL, SCJP, ACP, MB; Funding Acquisition, SCJP, ACP, MB.

## EXTENDED DATA

**Extended Data Figure 1 (related to Fig. 1).**
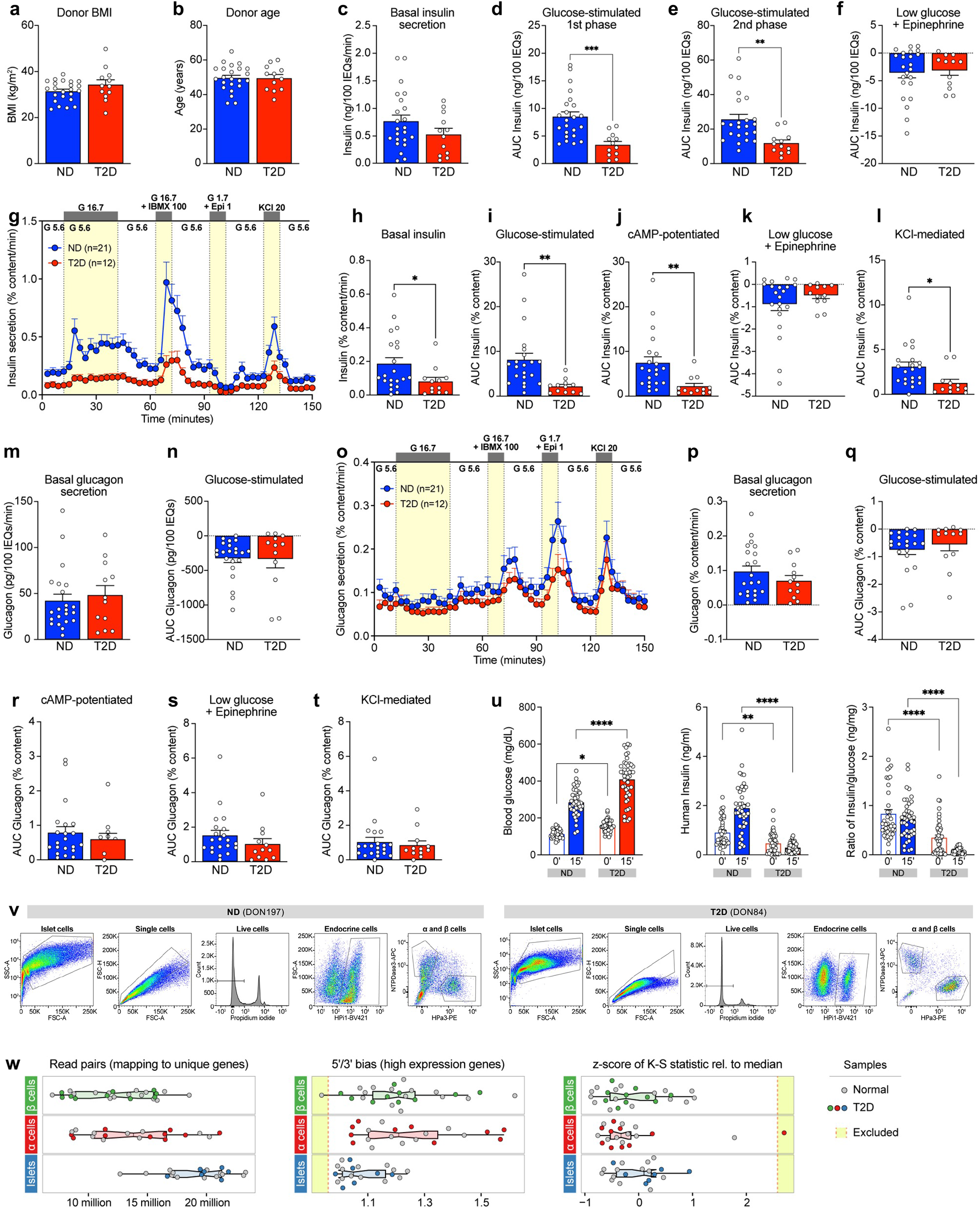
Additional metrics from functional and transcriptional profiling of islets from donors with short-duration T2D. **(a-b)** Matching of ND and T2D donor BMI **(a)** and age **(b)** for perifusion experiments. **(c)** Basal insulin secretion calculated as the average of the first three points of perifusion trace. **(d-e)** Integrated area under the curve **(AUC)** breaking down the total 16.7 mM glucose response into the first phase (**d**; through minute 24) and second phase (**e**; remainder of stimulation). **(f)** Area “under” the curve calculated from trace baseline for inhibition with low glucose and epinephrine. **(g-l)** Dynamic insulin secretion and metrics equivalent to Fig. 1 but normalized by total insulin content. **(m)** Basal glucagon secretion calculated as average of first three points of perifusion trace. **(n)** Area “under” the curve calculated from trace baseline for inhibition with high glucose. (**o-t**) Dynamic glucagon secretion and metrics equivalent to Fig. 1 but normalized by total glucagon content. **(u)** Blood glucose, human insulin levels, and human insulin:blood glucose ratio measured at 0’ (six-hour fasted) and 15’ after glucose and arginine stimulation of mice with human islet grafts. Symbols represent individual mice. * p<0.05, ** p<0.01, *** p<0.001, **** p<0.0001 (two-tailed t-test, panels **a-f**, **h-n**, and **p-t**; two-way ANOVA, panel **u**); error bars are SEM. **(v)** Gating strategy for sorted α and β cells identified by cell surface markers. Cell debris were excluded by forward scatter (FSC) and side scatter (SSC), single cells were identified by voltage pulse geometry (FSC-A v. FSC-H), and non-viable cells were excluded using propidium iodide (PI). Endocrine cell subpopulations were then gated based on positivity for HPi1 (pan-endocrine marker) and additional positivity for HPa3 (α cells) or NTPDase3 (β cells). **(w)** Select metrics used to assess library quality, organized by sample type. Outlier samples are highlighted in yellow and were excluded from downstream analyses.

**Extended Data Figure 2 (related to Fig. 1).**
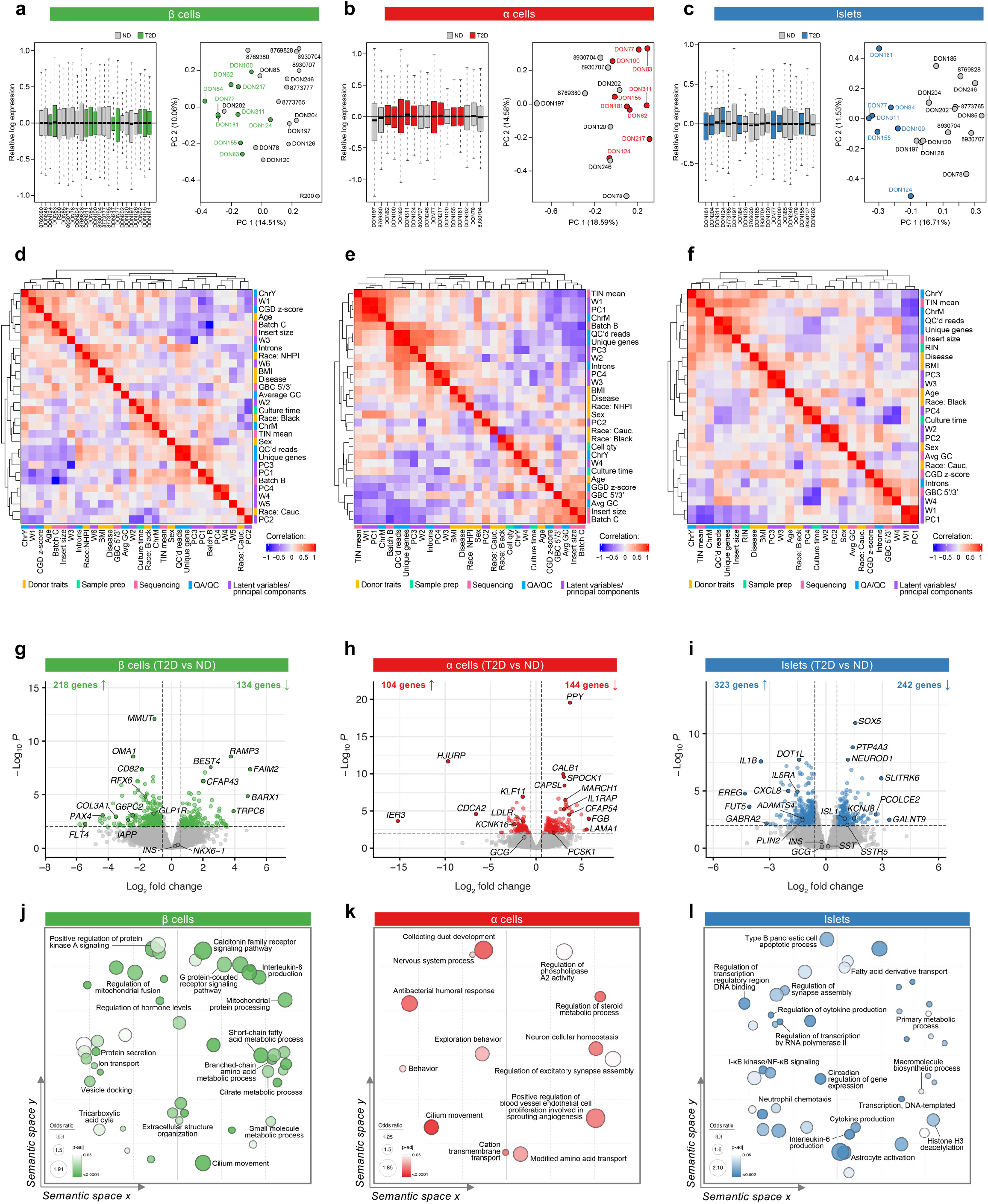
Transcriptional analysis of islets and sorted α and β cells reveals dysregulation of metabolic pathways in T2D β cells and immune signaling in T2D islets. (**a-c**) Relative expression of individual libraries post-correction and principal component (PC) analysis of each sample type. RRIDs (donor labels beginning with ‘8’) are abbreviated; see Extended Data Table 1 for complete alphanumeric RRIDs. Nondiabetic (ND) samples, grey; T2D samples, colored according to sample type. (**d-f**) Pearson correlation between sample covariates and PCs using the DEseq model. Colored bands next to row/column labels indicate whether variable is a donor trait (yellow), sample preparation variable (mint green), sequencing metric (pink), quality assurance/quality control (QA/QC) metric (blue), or latent variable or PC (purple). Culture time, duration of time (hours) between islet isolation and cell dispersion/sorting; Cell qty, number of sorted cells from which RNA was isolated (β and α cells only); RIN, RNA integrity number; Batch *x*, sequencing batch; TIN mean, mean transcript integrity number; Insert size, median length of sequenced RNA fragments; GBC 5’/3’, ratio of gene body coverage at 5’ and 3’ end, describing reads distribution along a gene; QC’d reads, number of read pairs that pass initial filters; unique reads, number of read pairs that map to genomic area covering exactly one gene; Introns, reads mapping to intronic regions of genes; Avg GC, average GC content of all reads; CGD z-score, z-score quantifying cumulative gene diversity of libraries from median based on Kolmogorov Smirnov test; ChrM, reads mapping to MT chromosome; ChrY, reads mapping to Y chromosome; PC*x*, principal components; W*x*, RUV-seq latent variables. (g-i) Volcano plots illustrating differentially expressed genes between ND and T2D β cells **(g)**, α cells **(h)**, and islets **(i)**. Lines denote cutoffs for fold-change (±1.5) and significance (<0.01); genes passing both thresholds are colored and select genes are labeled. (**j-l**) Enriched gene ontology terms (FDR<0.05) obtained from RNA Enrich were condensed using the RelSim function of Revigo (similarity=0.5) and plotted in semantic space to emphasize relatedness. Dot size represents odds ratio and color represents p-value. Select terms are labeled.

**Extended Data Figure 3 (related to Fig. 2).**
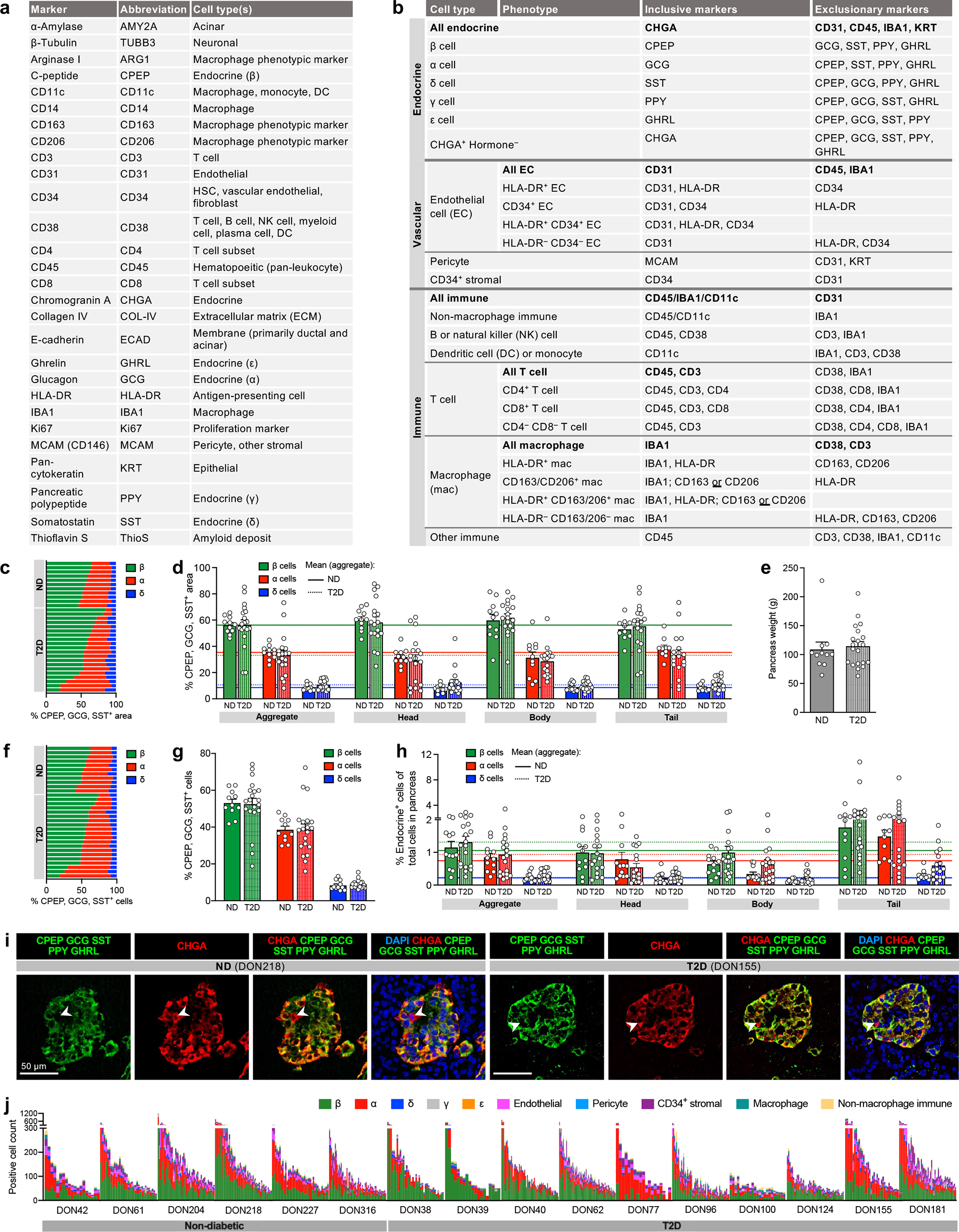
Parallel approaches of multiplexed imaging and high-throughput traditional immunohistochemistry enable profiling of endocrine cells in addition to intraislet vascular and immune cells. **(a-b)** Markers, cell populations, and specific phenotypes distinguished by the CODEX antibody panel. **(c-h)** Cross-sectional area (**c-d**) and cytonuclear quantification **(f-h)** of β cells (CPEP; green), α cells (GCG; red), and δ cells (SST; blue). Individual donor data shown in stacked bar graphs (**c,f**); bar graphs (**d-e, g-h**) show mean + SEM, one symbol per donor. Stratification by pancreas region (**d, g**) includes horizontal lines (solid, ND; dotted, T2D) for mean values from combined analysis (‘Aggregate). **(e)** Pancreas weight measured during organ procurement; used to calculate endocrine cell mass in **Fig. 2b**. **(i)** Representative images depicting rare cells positive for chromogranin A (CHGA; red) but negative for all hormones (green). Scale bars, 50 μm; arrowheads denote CHGA^+^ hormone^−^ cells. **(j)** Abundance of endocrine and non-endocrine cells in ND and T2D islets; one vertical bar per islet and colored by cell type. Islets are grouped by donor and ordered from largest (highest total cell number) to smallest. See also **Fig. 2e**.

**Extended Data Figure 4 (related to Fig. 2).**
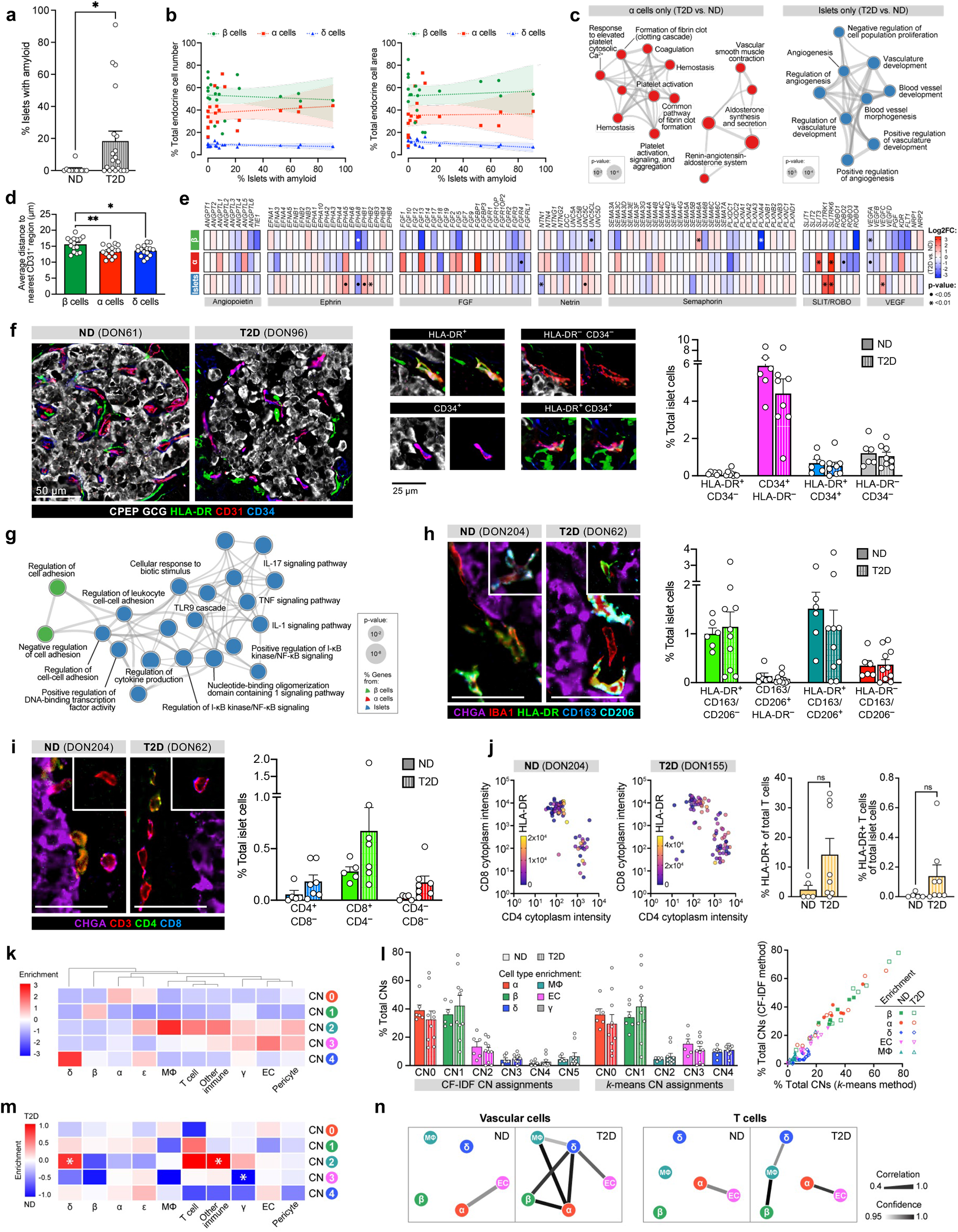
Integration of multiplexed imaging and transcriptional profiling highlight disrupted capillaries and immune cells within T2D islets. **(a)** Amyloid prevalence (% total islets with amyloid, averaged over multiple regions); * p<0.05 (two-tailed t-test). **(b)** Correlation of amyloid prevalence with β, α, and δ cell populations as percentage of total endocrine cell number or cross-sectional area; one symbol per donor with 95% confidence interval of linear regression (shading). No slopes were significantly nonzero at p<0.01 threshold. **(c)** Metascape visualization of select terms enriched for differentially expressed genes in T2D α cells (left) and islets (right). **(d)** Average distance of each endocrine cell type to nearest capillary; one symbol per donor (both ND and T2D); asterisks signify results of one-way ANOVA with Tukey’s multiple comparisons test (** p<0.01; * p<0.05). **(e)** Gene expression fold-change of selected vascular and neuronal ligands and their receptors in β cells, α cells, and islets; • FDR<0.05; * FDR<0.01. **(f)** Phenotypes of endothelial cells (CD31; red) defined by single or dual positivity for HLA-DR (green) and CD34 (blue). Examples of each combination (HLA-DR^+^ CD34^−^, CD34^+^ HLA-DR^−^, HLA-DR^+^ CD34^+^, and HLA-DR^−^ CD34^−^) are shown to right. **(g)** Magnification of select clusters depicted in **Fig. 1r** (terms enriched across β, α, and islet samples). (**h-i**) Macrophages (IBA1^+^) and T cells (CD3^+^) phenotyped by various cell surface markers; insets show additional cells to illustrate phenotypic variety. Scale bars, 50 μm. **(j)** Expression of HLA-DR in CD4^+^ and CD8^+^ T cell populations. **(k-l)** Cellular neighborhood assignment and corresponding cell composition changes in T2D vs. ND islets. Panels **k** and **m-n** show results from the *k*-means method and panel **l** compares these results to CF-IDF method shown in **Fig. 2o-2s**. Traditional IHC data: panels **a-b**; CODEX data: panels **d**, **f**, **h-n**. RNA data: panels **c**, **e**, **g**.

**Extended Data Figure 5 (related to Fig. 3).**
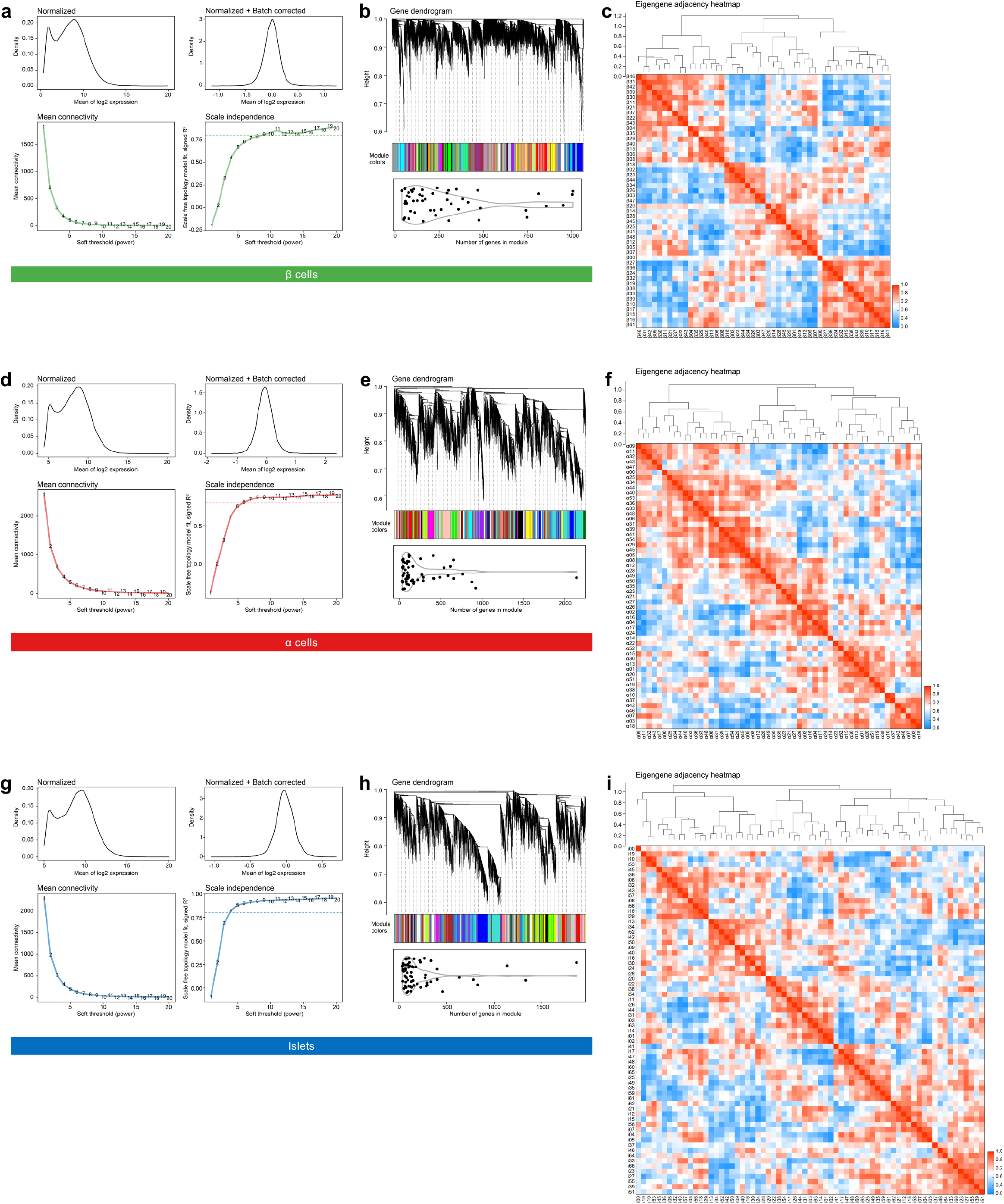
Quality assessment of Weighted Gene Co-expression Network Analysis (WGCNA). Analyses for β cell **(a-c)**, α cell **(d-f)**, and islet **(g-i)** datasets were conducted in parallel. Metrics are shown for batch correction and network parameter selection (**a, d, g**), module size and assignment (**b, e, h**), and module relatedness (**c, f, i**).

**Extended Data Figure 6 (related to Fig. 3).**
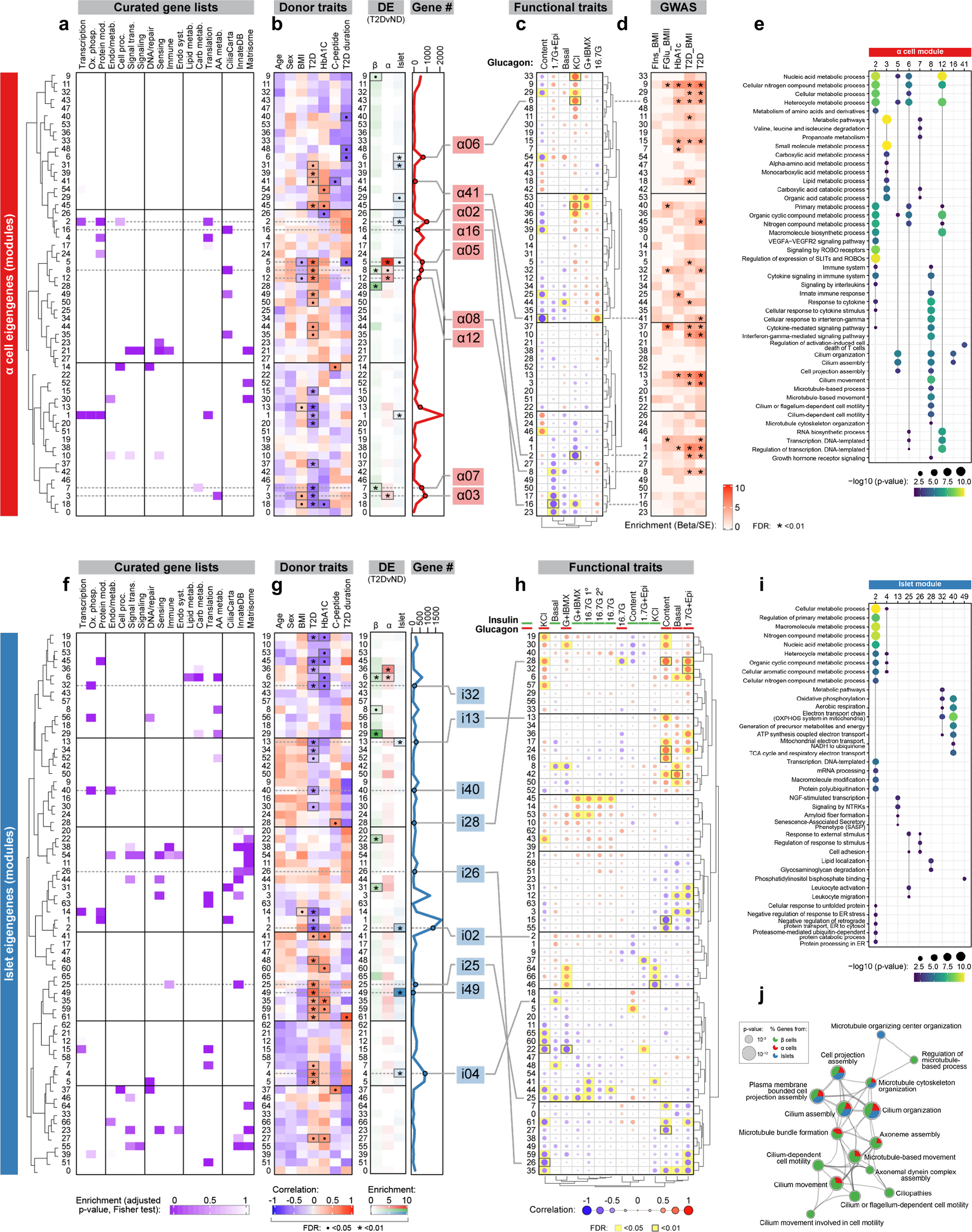
WGCNA emphasizes α and islet cell gene modules associated with donor and islet traits as well as those enriched in GWAS loci. Module eigengenes for α cells **(a-e)** and islets **(f-i)** shown in parallel to β cells (**Fig. 3a-3e**). (**a, f**) Modules clustered by similarity and showing relative enrichment of curated gene lists. (**b, g**) Module correlation to donor characteristics, enrichment of differentially expressed (DE) genes, and total number of genes per module. • p<0.05; * p<0.01. Modules of interest highlighted (b: red, g: blue). (c, h) Module correlation to α and β cell function (Fig. 1); significant associations highlighted (yellow). For islets **(g)**, modules were correlated to both insulin and glucagon secretion. G+IBMX, 16.7 mM glucose with 100 μM isobutylmethylxanthine; 16.7G, 16.7 mM glucose; 16.7G 1°, first phase; 16.7G 2°, second phase; 1.7G+Epi, 1.7 mM glucose and 1μM epinephrine; KCl, 20 mM potassium chloride. **(d)** Module enrichment for GWAS traits. FIns, fasting insulin; FGlu, fasting glucose. * FDR<0.01. (e, i) Enrichment of select gene ontology terms in β cell modules with notable correlations and/or enrichment. **(j)** Magnification of select clusters depicted in Fig. 1r (terms enriched across β, α, and islet samples).

**Extended Data Figure 7 (related to Fig. 4).**
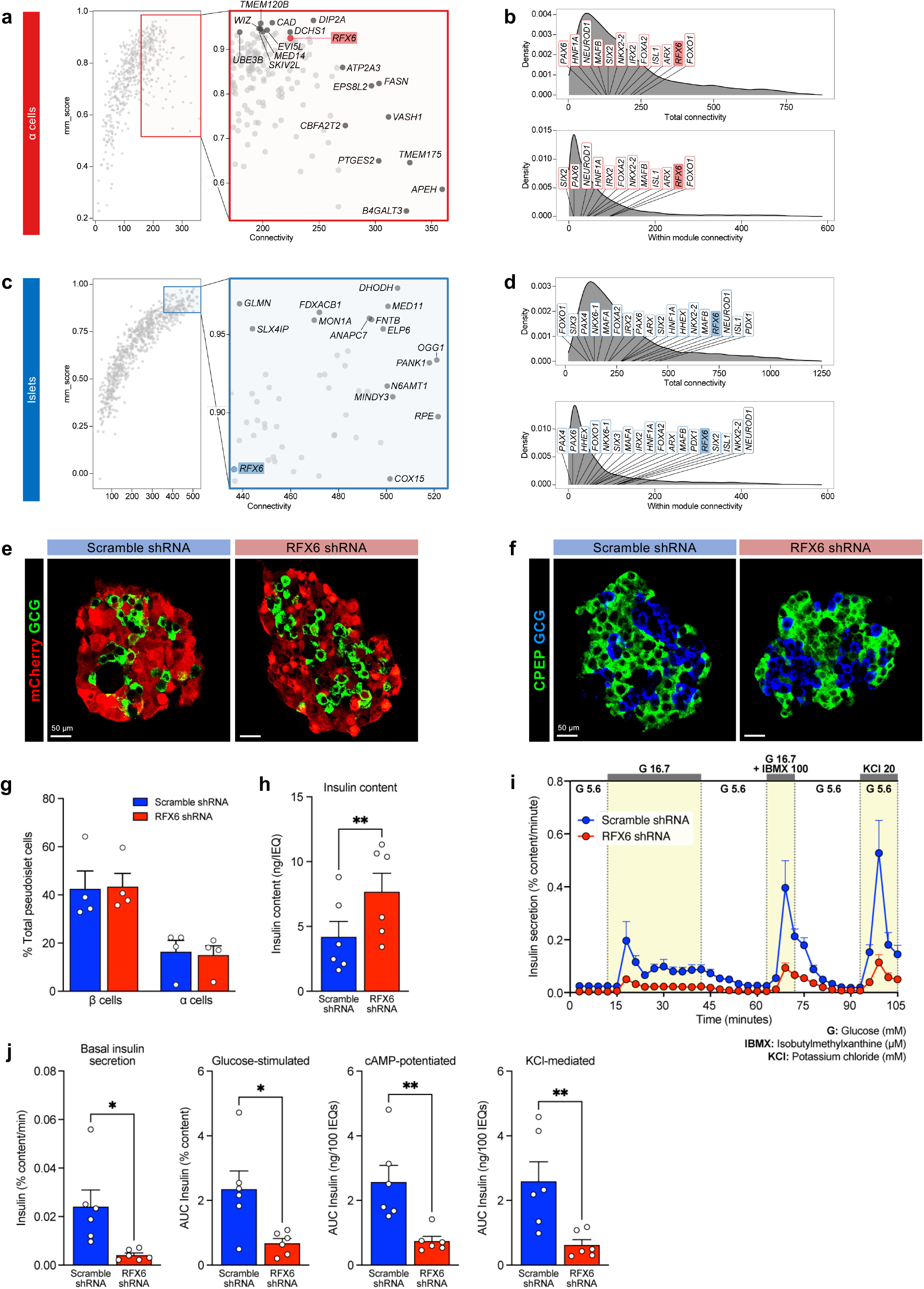
Connectivity of *RFX6* by WGCNA is β cell-specific and *RFX6* reduction impairs insulin secretion. **(a-d)** Connectivity of genes in α cell **(a-b)** and islet **(c-d)** modules, in parallel to data for β cell modules in Fig. 4a-4b. **(a, c)** Overall connectivity of individual genes; select genes with high connectivity scores are labeled. **(b, d)** Cross-and within-module connectivity; select transcription factors are labeled. **(e-f)** Immunofluorescent staining of pseudoislets embedded in type I collagen. **(e)** Transduced α cells marked by mCherry; see Fig. 4h for β cells. **(f)** Distribution of β cells (CPEP; green) and α cells (GCG; blue). **(g)** Quantification of % β and % α cells in control (scramble) and sh*RFX6* pseudoislets. **(h)** Insulin content in control and sh*RFX6* pseudoislets (** p<0.01, two-tailed t-test). **(i-j)** Dynamic insulin secretion and metrics equivalent to Fig. 4j-4k but normalized by total insulin content.

**Extended Data Figure 8 (related to Fig. 5).**
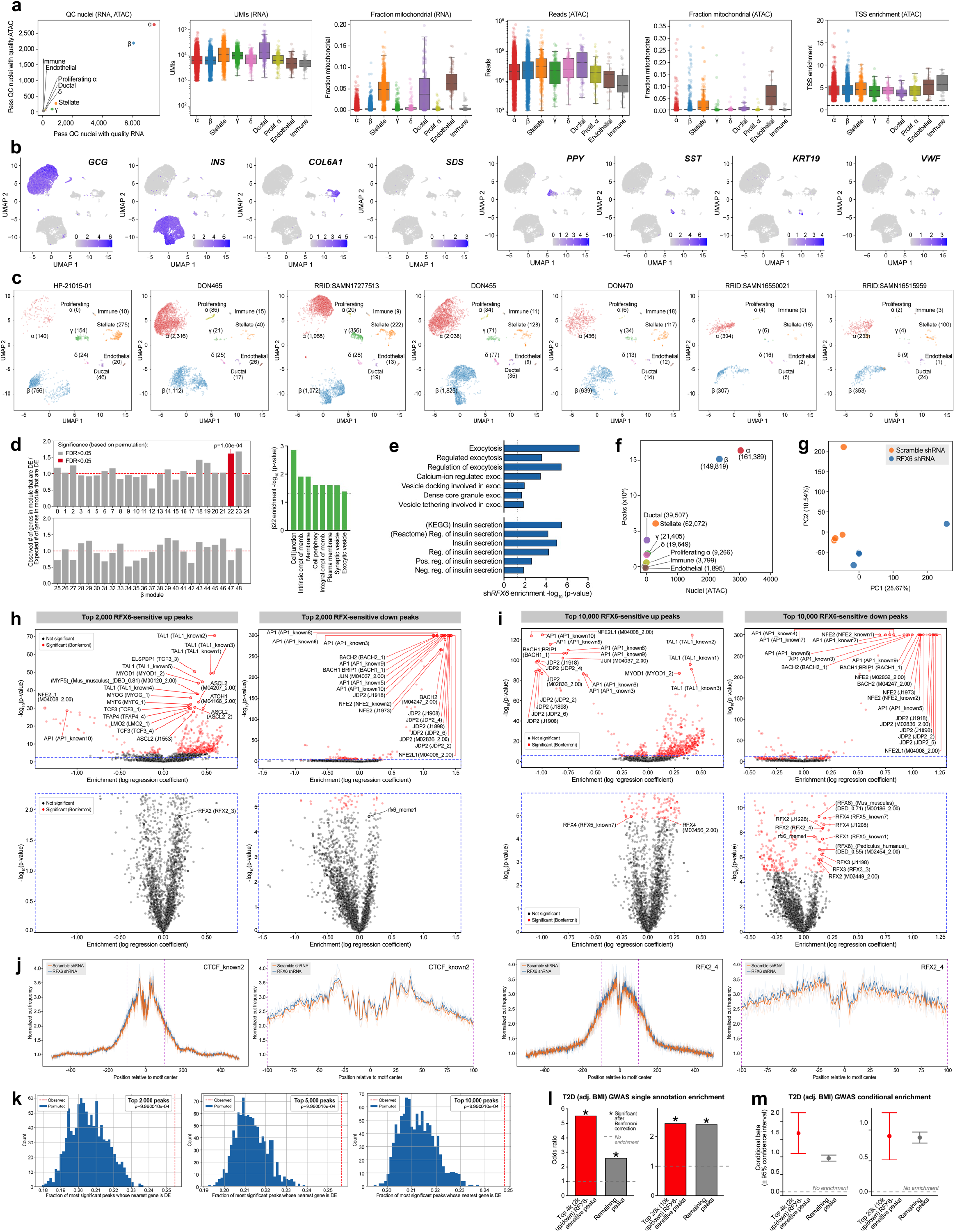
Application of dual RNA and ATAC-sequencing to single nuclei from *RFX6* shRNA pseudoislets. **(a)** Quality control of nuclei for RNA and ATAC modalities. UMI, unique molecular identifier; TSS, transcription start site. **(b)** Expression of marker genes in cell type clusters. **(c)** Per-donor cell type counts. See also Fig. 5b-5c. **(d)** Enrichment of sh*RFX6* β cell nuclei differentially expressed genes within each β cell module derived from transcriptomes of sorted ND and T2D β cells (see Fig. 3a-3d). Right panel: cell component (cmpt) terms enriched for genes in β module 22. Memb., membrane. **(e)** Membership enrichment for exocytosis (exoc.) and insulin secretory pathways based on sh*RFX6* β cell nuclei differentially expressed (p<0.01) genes. All pathways are GO terms unless otherwise indicated. Neg., negative; pos., reg., regulation. **(f)** Per-cluster ATAC peaks (exact number listed in parentheses next to cell type). **(g)** PCA of pseudobulk β cell ATAC peak signal, each marker representing nuclei from a single donor/construct combination. (h-i) Motif enrichment for top 2,000 **(h)** or 10,000 **(i)** RFX6-sensitive up- and downregulated ATAC peaks in sh*RFX6* β cell nuclei. Motifs with highest significance are labeled in top panels; significant RFX motifs (or the single RFX motif closest to significance, in the case that no RFX motifs reach significance) are labeled in bottom panels. **(j)** ATAC footprints for CTCF_known2 and RFX2_4 motifs in β cell ATAC peaks. Light lines represent per-donor footprints; bold lines represent the average across donors. **(k)** Enrichment of top RFX6-sensitive up- and downregulated ATAC peaks (n=2,000, 5,000, or 10,000) in sh*RFX6* β cell nuclei near sh*RFX6* β cell differentially expressed genes. (**l-m**) Odds ratio of T2D GWAS enrichment **(l)** and model estimate from conditional analysis **(m)** of top 2,000 or 10,000 RFX6-sensitive peaks.

**Extended Data Figure 9 (related to Fig. 5).**
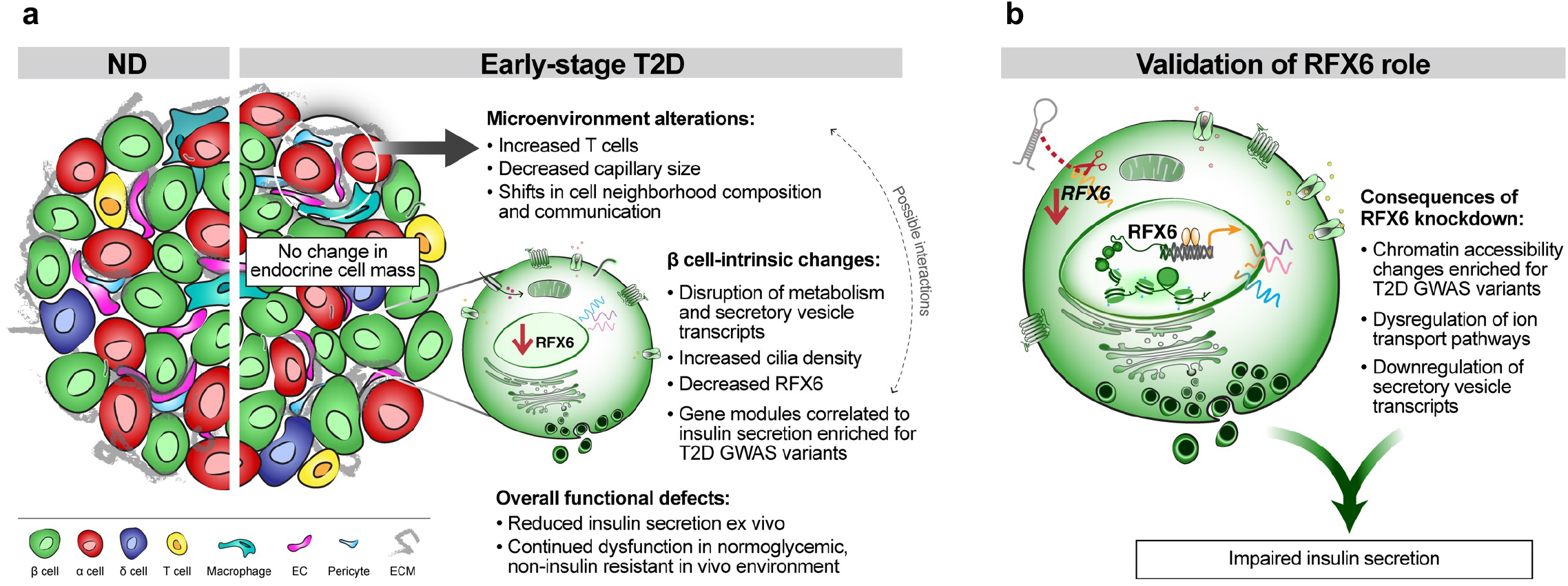
RFX6-mediated chromatin, transcriptome, and insulin secretion dysregulation in human β cells. **(a)** Major β cell-intrinsic and islet microenvironment alterations that define islet dysfunction in early-stage T2D. Observations from transcriptomic and histologic studies revealed no change to endocrine cell composition but evidence of dysregulated β cell processes and modest changes to intraislet vascular and immune cell populations. Insulin secretion was reduced and persisted in a nondiabetic environment. **(b)** RFX6 knockdown using a primary human pseudoislet system resulted in dysregulated vesicle trafficking and ion transport pathways, mediated by chromatin architectural changes overlapping with T2D GWAS variants. This led to reduced insulin secretion, confirming the critical role of RFX6 in human β cell function.

**Extended Data Table 1.**
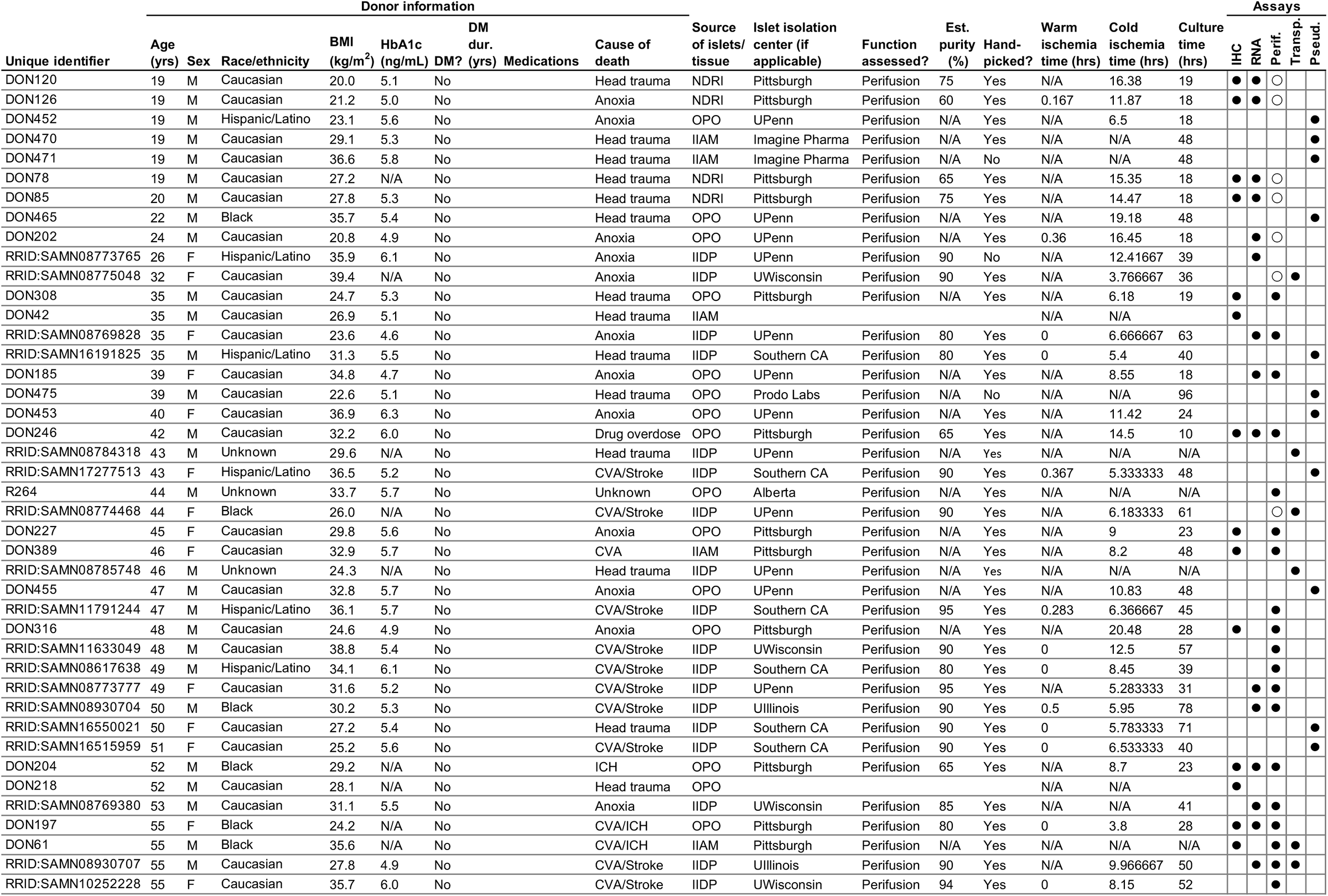
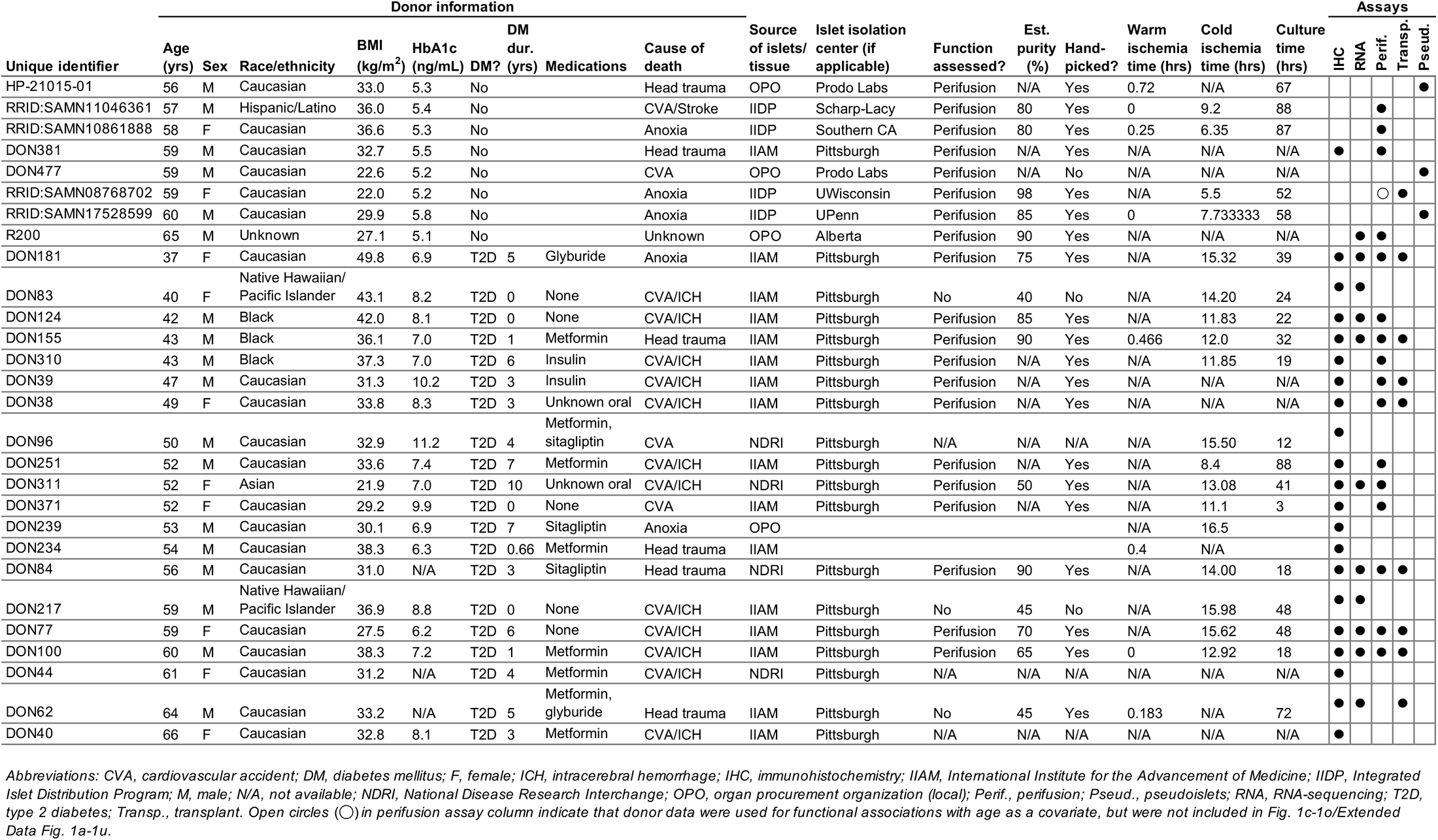
Donor characteristics, sample types, and experimental usage. Adapted from Hart NJ, Powers AC (2018), Checklist for Reporting Human Islet Preparations Used in Research, Diabetologia, doi.org/10.1007/s00125-018-4772-2.

**Extended Data Table 2.**
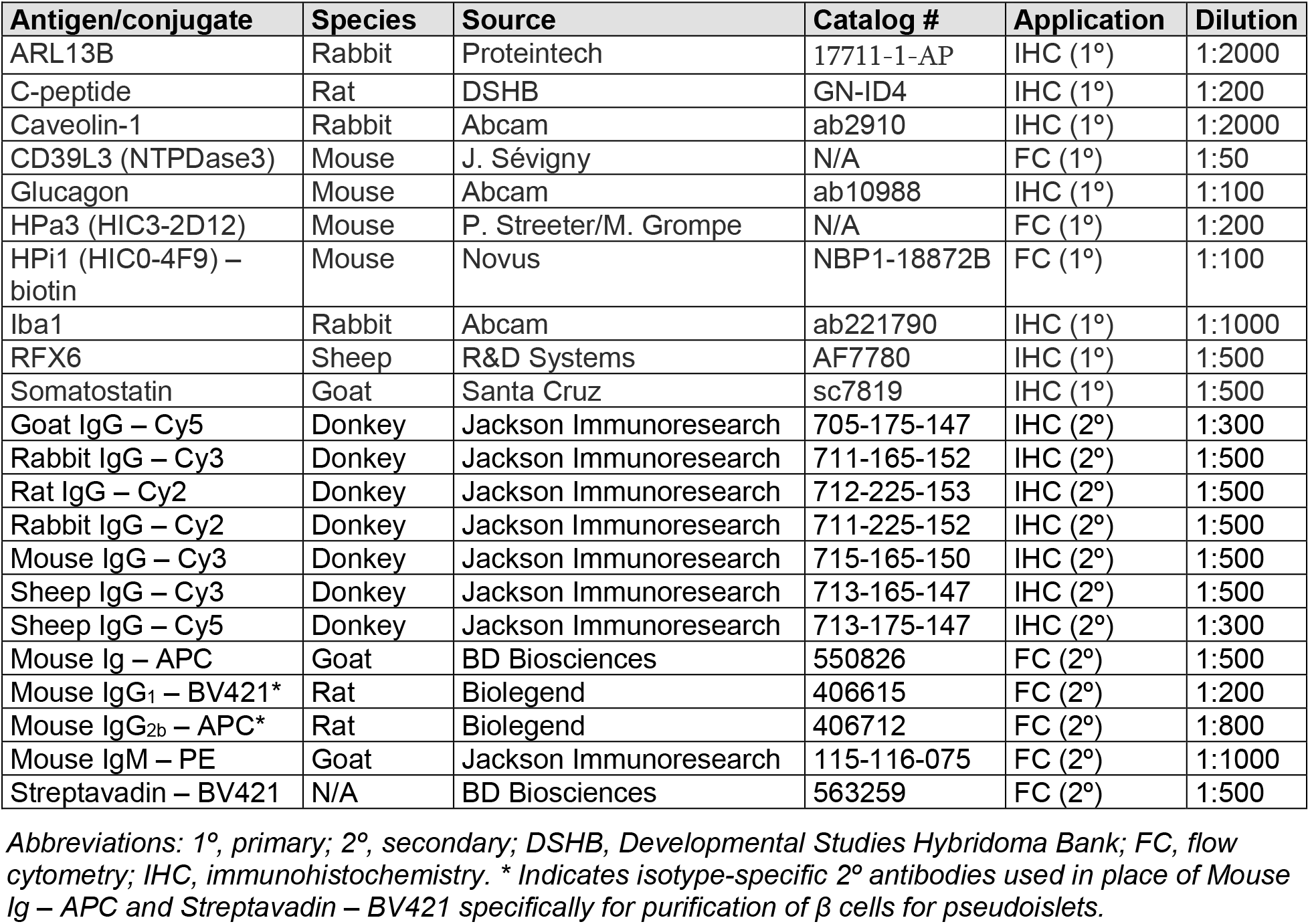
Antibodies for traditional immunohistochemistry and flow cytometry.

**Extended Data Table 3.**
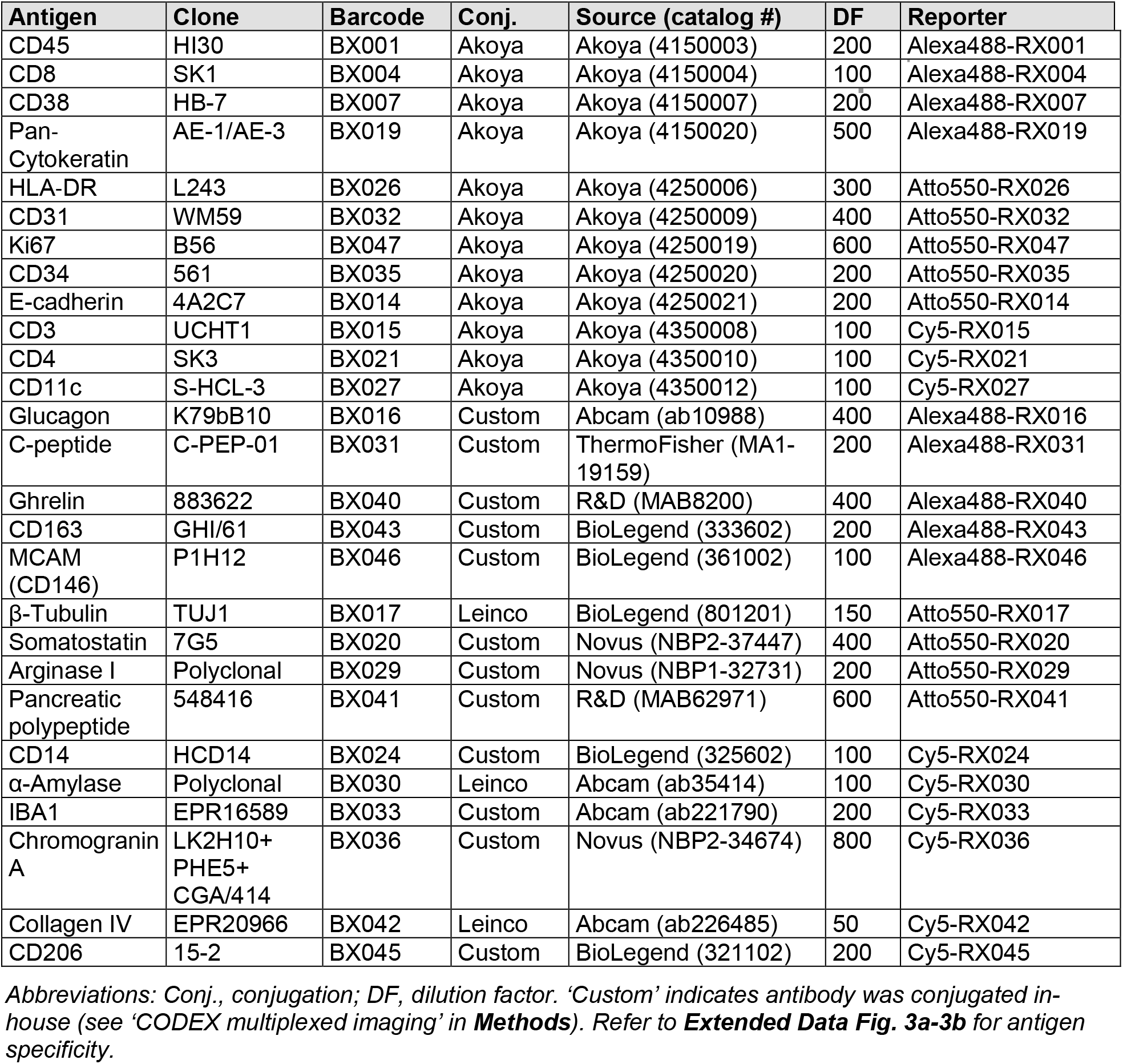
Antibodies for co-detection by indexing (CODEX).

**Extended Data Table 4.** Curated gene lists for weighted gene co-expression network analysis.

| Abbreviation | Full name | Source | List(s) |
| --- | --- | --- | --- |
| Transcription | -- | KEGG Pathway<sup>a</sup> | 2.1 |
| Ox. phosp. | Oxidative phosphorylation | KEGG Pathway<sup>a</sup> | 1.2 (00190 only) |
| Protein mod. | Folding, sorting and degradation | KEGG Pathway<sup>a</sup> | 2.3 |
| Endo/metab. | Endocrine and metabolic disease | KEGG Pathway<sup>a</sup> | 6.10 |
| Cell proc. | Cell growth and death | KEGG Pathway<sup>a</sup> | 4.2 (04110, 04210, 04216, 04217, 04115, 04218 only) |
| Signal trans. | Signal transduction | KEGG Pathway<sup>a</sup> | 3.2 |
| DNA/repair | Replication and repair | KEGG Pathway<sup>a</sup> | 2.4 |
| Sensing | Signaling molecules and interaction | KEGG Pathway<sup>a</sup> | 3.3 |
| Immune | Immune system | KEGG Pathway<sup>a</sup> | 5.1 |
| Endo syst. | Endocrine system | KEGG Pathway<sup>a</sup> | 5.2 |
| Lipid metab. | Lipid metabolism | KEGG Pathway<sup>a</sup> | 1.3 |
| Carb metab. | Carbohydrate metabolism | KEGG Pathway<sup>a</sup> | 1.1 |
| Translation | -- | KEGG Pathway<sup>a</sup> | 2.2 |
| AA metab. | Amino acid metabolism | KEGG Pathway<sup>a</sup> | 1.5, 1.6 |
| CiliaCarta<sup>b</sup> | -- | van Dam et al. 2019<sup>32</sup> | -- |
| InnateDB<sup>c</sup> | -- | Breuer et al. 2013<sup>31</sup> | Immunome Database |
| Matrisome | -- | GSEA<sup>d</sup> | M5889<sup>30</sup> |

<sup>a</sup>Available https://www.genome.jp/kegg/pathway.html
<sup>b</sup>Available http://bioinformatics.bio.uu.nl/john/syscilia/ciliacarta/
<sup>c</sup>Available https://www.innatedb.com/redirect.do?go=resourcesGeneLists
<sup>d</sup>Available https://www.gsea-msigdb.org/gsea/msigdb/genesets.jsp

